# Septin-associated PIPKIγ splice variants drive centralspindlin association with the midbody via PI(4,5)P_2_

**DOI:** 10.1101/2024.02.22.581562

**Authors:** Giulia Russo, Nadja Hümpfer, Nina Jaensch, Steffen Restel, Christopher Schmied, Florian Heyd, Martin Lehmann, Volker Haucke, Helge Ewers, Michael Krauss

## Abstract

Mammalian cytokinesis critically depends on the phospholipid phosphatidylinositol 4,5-bisphosphate [PI(4,5)P_2_] which serves as docking site for crucial components of the cytokinetic machinery at the plasma membrane. PI(4,5)P_2_ supports several stages of cytokinesis, including actomyosin ring assembly and constriction, membrane tethering of spindle microtubules, and midbody organization. How these various activities of PI(4,5)P_2_ and the underlying mechanisms of local PI(4,5)P_2_ synthesis are orchestrated in space and time has remained elusive. Here, we identify a pivotal role of septin-binding splice variants of PIPKIγ that couple nanoscale PI(4,5)P_2_ synthesis at the ingressing cleavage furrow to late midbody formation. Depletion of PIPKIγ isoforms causes multinucleation and perturbs anillin and septin deposition at the intercellular bridge and at the midbody. These defects are rescued by wild-type kinase, but not by septin binding-deficient or catalytically inactive PIPKIγ variants. We further show that both, septins and PIPKIγ form a complex with centralspindlin, and thereby facilitate the recruitment of centralspindlin to the midbody. Taken together, our findings establish septin-associated PIPKIγ isoforms as key regulators of midbody organization that act through generating a local pool of PI4,5P_2_ required for centralspindlin recruitment and maintenance at the midbody, and for septin association with microtubules.

## Introduction

Cytokinesis is the final step of cell division that ultimately partitions the cytoplasmic content of the mother cell into two daughter cells. Failures in cytokinesis can cause tetraploidy, centrosome amplification and chromosomal instability, and may promote tumorigenesis^1^. Cytokinesis starts with the assembly of a contractile actomyosin ring at the equatorial plane of the mother cell that drives the formation of the cleavage furrow^2,3^. The cleavage furrow ingresses symmetrically, until the daughter cells are connected only by a thin intercellular bridge (ICB) formed around a tight bundle of central spindle microtubules^4^, the cytokinetic bridge. As the ICB elongates, a dense structure assembles and matures in its center, the midbody, an organelle of complex and unique protein and lipid composition^5^. The midbody serves several important functions, including tethering of spindle microtubules to the plasma membrane, and the orchestration of the abscission machinery^6^.

Cytokinesis is accompanied by drastic morphological changes of the mother cell, which rely on constant remodeling and reorganization of the cytoskeleton. These events are concerted by phosphoinositides^7^, in particular by the plasma membrane-enriched phosphatidylinositol 4,5-bisphosphate [PI(4,5)P_2_]^8^. PI(4,5)P_2_ serves as a docking site for several components of the cytokinetic machinery^9^. It accumulates early at the newly forming cleavage furrow, where it anchors the contractile machinery to the cell equator to ensure symmetric ingression of the furrow^10^. This process is orchestrated by anillin, a multidomain protein that scaffolds actin, myosin, septins and the central spindle^11^. Through a carboxyterminal PI(4,5)P_2_-binding pleckstrin homology (PH) anillin attaches this scaffold to the plasma membrane^12,13^.

As the contractile ring closes PI(4,5)P_2_ further concentrates along the cleavage furrow, and later at the ICB to support bridge stabilization^10,14^. ICB stability likewise depends on the presence of septins, a family of filament-forming, GTP-binding proteins^15,16^ that are recruited to the cleavage furrow through a direct interaction with anillin^10,12^. Septins are required for symmetric cleavage furrow ingression, and by an unknown mechanism appear to promote stable anchorage of the ingressed furrow at the cytokinetic bridge, to prevent furrow regression upon completion of contraction^17,18^. Septin filaments associate with membranes enriched in PI(4,5)P_2_ in vitro and in living cells^19–21^, and several lines of evidence indicate that PI(4,5)_2_ is required for the localization of septins at the ICB (reviewed in ^22^).

At later stages of cytokinesis, when the midbody has formed, PI(4,5)P_2_ tethers spindle microtubules to the plasma membrane through centralspindlin^23^, a constitutive heterotetramer of MKLP1 (also known as KIF23) and MgcRacGAP (also known as RacGAP1)^24^. Whereas MKLP1 promotes bundling of antiparallel spindle microtubules at the midbody, MgcRacGAP locally controls the activities of Rho GTPases^25^ and through an atypical C1 domain associates with plasmalemmal PI(4,5)P_2_ and PI(4)P^23^. At the same time, PI(4,5)P_2_ facilitates exocyst docking to the plasma membrane of the ICB, and thereby supports the abscission process^26^.

Though clearly essential at multiple stages of cytokinesis it has remained an important, yet unresolved question how PI(4,5)P_2_ synthesis is mechanistically coupled to cytokinetic progression. In mammalian cells PI(4,5)P_2_ is predominantly generated by type I PI(4)P 5-kinases, which are encoded by three genes named PIPKIα, Iβ and Iγ, with several splice isoforms being known for PIPKIγ^27^. All isozymes are widely expressed in different tissues and cell types, but they exhibit unique subcellular distributions, and trigger the formation of functionally distinct pools of PI(4,5)P_2_. Mechanisms underlying the specific subcellular distribution of select isozymes are intensely investigated. In case of PIPKIγ short splice inserts target kinase activity to distinct locations for a variety of subcellular events. Isoform i2 for instance contains a short peptide that associates with AP-2 and talin, to promote kinase recruitment to sites of endocytosis^28–30^ and to focal adhesions^31,32^, respectively. Isoform i5 populates endosomal compartments^33^ and autophagosomes^34^, due to its interactions with SNX5 and Atg14. PIPKIγ recruitment factors are are often also PI(4,5)P_2_ effectors, and, thus, considered to be part of a feed-forward loop that enhances PI(4,5)P_2_ synthesis in their immediate proximity to reinforce their own membrane association^35^. It is likely that similar mechanisms underly the recruitment and tethering of cytokinetic proteins, but up to now no interactions with the PI(4,5)P_2_-synthesizing machinery have been revealed.

Here, we report a novel functional role of PIPKIγ during ICB maturation and late midbody formation. We demonstrate that splice isoforms i3 and i5 associate with septins, and that this interaction is essential for the retention of anillin and septins at the ICB. Our analyses further demonstrate that the septin-dependent recruitment of PIPKIγ to the ICB is required for efficient targeting of centralspindlin to the midbody, and for the translocation of septins onto microtubules at late stages of cytokinesis.

## Results

### PIPKIγ is required at early and late stages of cytokinesis

The pivotal role of PI(4,5)P_2_ during cytokinesis is well established, but little is known about the enzymes mediating its synthesis at cytokinetic stages. In mammalian cells PI(4,5)P_2_ is mainly synthesized by type I PIPK enzymes, named PIPKIα, Iβ and Iγ. We depleted individual isozymes in HeLa cells, and subsequently analyzed the cells for cytokinetic defects. To this end we immunostained for acetylated tubulin, a marker of spindle microtubules before furrow ingression, and of microtubules within the ICB from telophase on until abscission (Fig. 1A). Approximately 2 % of the cells displayed a mitotic spindle under control conditions (Fig. 1B, Fig. S1A/B). Depletion of PIPKIβ, but not of PIPKIα significantly increased this fraction, in line with previous reports^14^. Likewise, loss of PIPKIγ increased the fraction of cells exhibiting an acetylated tubulin bridge, i.e. cells in meta- and anaphase. This was surprising as this isozyme is believed to contribute only a minor pool of PI(4,5)P_2_ in non-neuronal cells. More importantly, however, depletion of PIPKIγ, but not of Iα or Iβ, doubled the fraction of cells exhibiting an acetylated tubulin bridge, indicating an arrest in telophase (Fig. 1C), and significantly increased the percentage of multi-nucleated cells about threefold (Fig. S1C/D).

**Figure 1:**
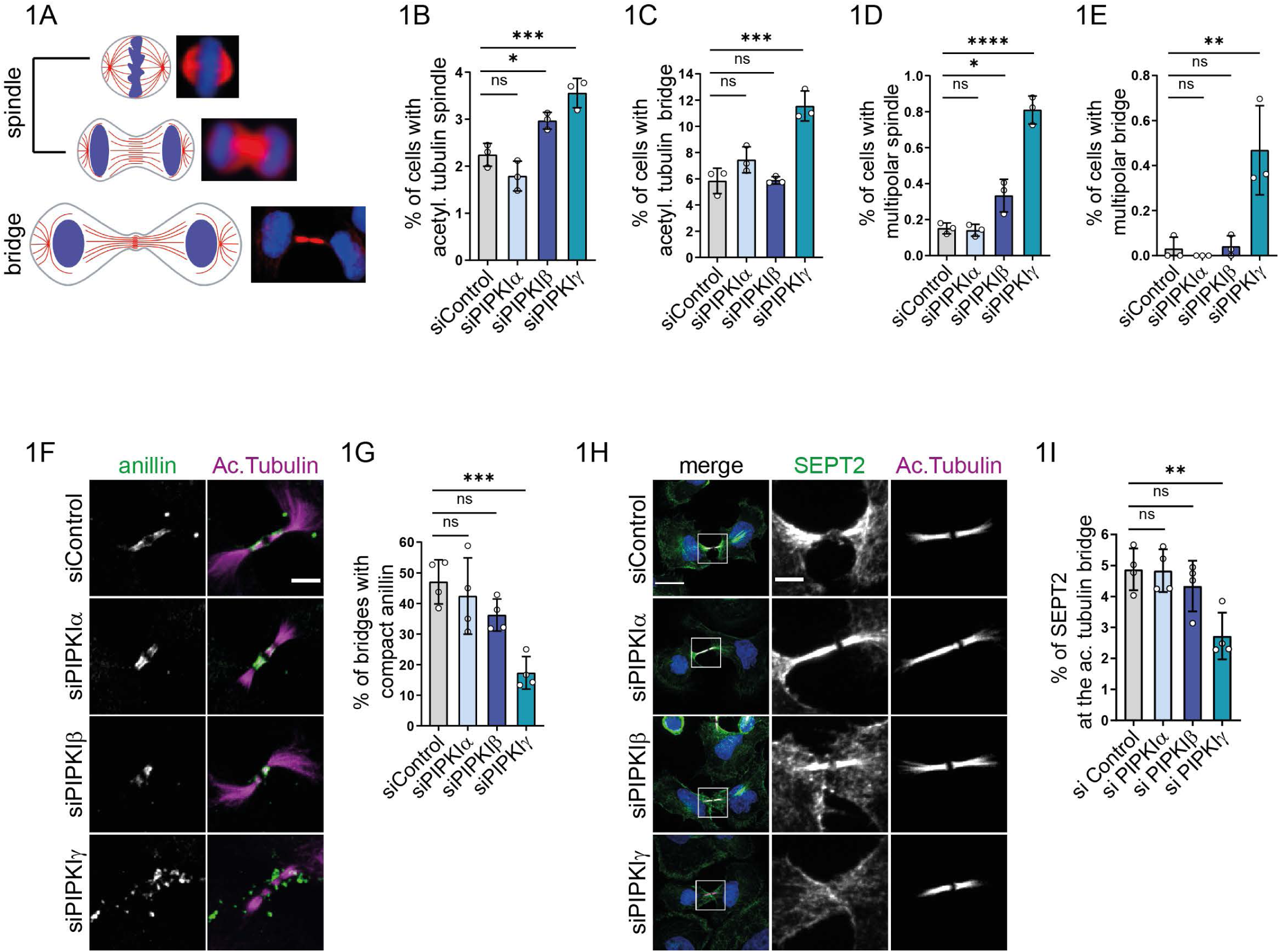
PIPKIγ is required for cytokinetic progression, and for the organization of anillin and septins at the ICB. **(A-E)** Depletion of PIPKIβ or of PIPKIγ stalls cells at early stages of mitosis, while exclusively PIPKIγ controls mitotic progression after furrow ingression. Cells were depleted of individual PIPKI isozymes, fixed and stained for acetylated tubulin, and analyzed by semi-automated imaging (see Figure S1B for exemplary images). Data are represented as mean ± SD (n=3, with 2593-4431 cells imaged per condition per experiment. Statistics: One-way ANOVA, followed by Dunnett’s multiple comparison test. **(A)** Left: Scheme depicting the organization of acetylated microtubules (red lines) in mitotic cells before furrow ingression, and upon formation of the ICB. Right: Representative images of mitotic cells upon staining for acetylated tubulin (in red). **(B)** Percentage of cells displaying a central spindle (adjusted P values: siControl vs. siPIPKIα, P=0.1723; siControl vs. siPIPKIβ, P=0.0274; siControl vs. siPIPKIγ, P=0.0009), or **(C)** bundled microtubules at the ICB (adjusted P values: siControl vs. siPIPKIα, P=0.1423; siControl vs. siPIPKIβ, P=0.0007; siControl vs. siPIPKIγ, P=0.0002). **(D)** Percentage of cells with multipolar spindles (adjusted P values: siControl vs. siPIPKIα, P=0.9933; siControl vs. siPIPKIβ, P=0.0203; siControl vs. siPIPKIγ, P<0.0001). or **I** multipolar ICBs (adjusted P values: siControl vs. siPIPKIα, P=0.9701; siControl vs siPIPKIβ, P=0.9986; siControl vs. siPIPKIγ, P=0.0024). **(F-G**) Depletion of PIPKIγ scatters anillin at the ICB. **(F)** Representative confocal images (max intensity z-projections) of HeLa cells treated with siRNA control, or against PIPKI α, β, or γ, synchronized at late cytokinesis, and stained for acetylated tubulin and anillin. Note that the brightness of acetylated tubulin needed to be slightly increased to be able to visualize its localization upon knockdown of PIPKIγ. Scale bar: 5μm. **(G)** Percentage of ICBs with compact anillin. Data are represented as mean ± SD. (n=4, 15-30 bridges were imaged per condition per experiment). Statistics: One-way ANOVA followed by Dunn’tt’s multiple comparison test (adjusted P values: siControl vs. siPIPKIα, P=0.7600; siControl vs. siPIPKIβ, P=0.1914; siControl vs. siPIPKIγ, P=0.0006). **(H-I)** Depletion of PIPKIγ decreases the fraction of SEPT2 localizing at the acetylated tubulin bridge. **(H)** Representative confocal images (max intensity z-projection) of HeLa treated with siRNA control or against PIPKI α, β,γ, synchronized at late cytokinesis and stained for acetylated tubulin and SEPT2. Scale bar of merge: 20μm, scale bar of grey insets: 5μm. **(I)** Percentage of total SEPT2 at the acetylated tubulin bridge. Quantification was performed on average intensity projections after background subtraction. SEPT2 intensity at the acetylated tubulin bridge was divided by total intensity of SEPT2 from dividing cell. Values are represented as mean ± SD (n=4; 15-30 cytokinetic cells were imaged per condition per experiment). Statistics: One-way ANOVA, followed by Dunn’tt’s multiple comparison test (adjusted P values: siControl vs. siPIPKIα, P=0.9996, siControl vs. siPIPKIβ, P=0.6167; siControl vs. siPIPKIγ, P=0.0038).

Our analyses also revealed abnormalities such as multipolar spindles or bridges. Again, depletion of PIPKIβ or Iγ caused a significant increase in the fraction of cells that displayed a multipolar spindle (Fig. 1D), while solely the depletion of PIPKIγ caused a significant increase in cells with multipolar bridges (Fig. 1E). Depletion of neither type I PIPK isozyme significantly changed total PI(4,5)P_2_ levels at the plasma membrane (Fig. S1E/F). This was expected, as despite their unique tissue distribution and distinct subcellular localizations PIPK Iα, Iβ, and Iγ are able to largely compensate for each other^36^.

Taken together, these data reveal that PIPKIβ and Iγ have synergistic roles during furrow ingression, while PIPKIγ serves a so far unknown, unique function at later stages of cytokinesis, when the ICB is formed.

### PIPKIγ is required to concentrate the PI(4,5)P_2_ effectors anillin and septins during ICB maturation

Given the unique defects observed upon depletion of PIPKIγ at late stages of cytokinesis, we analyzed the subcellular distribution of PI(4,5)P_2_ effectors in HeLa cells synchronized at the stage where they exhibit a mature ICB. Anillin bears a PI(4,5)P_2_-binding PH domain in its C-terminus^11,12^ that facilitates its recruitment to the equatorial plane of the dividing cell. At cytokinesis, prior to abscission, anillin is found concentrated at the midbody, and at its flanks^37^ (Fig. 1F). Control cells, as well as cells lacking expression of PIPKIα or Iβ, displayed a compact organization of anillin at the ICB, as visualized by co-staining of acetylated tubulin. In absence of PIPKIγ, however, anillin was found scattered along the bridge. Additional clusters of anillin were detected in areas adjacent to the bridge, which were rarely observed under control conditions. As a result, the fraction of cells displaying compact anillin significantly declined in PIPKIγ knockdown cells (Fig. 1G).

Through its C-terminal PH domain anillin also associates with septins^38^, a family of guanine nucleotide-binding proteins that hetero-oligomerize into hexamers and octamers. PI(4,5)P_2_ promotes their association with membrane surfaces in living cells and in vitro^19–21^. Though largely dispensable during furrow ingression in mammalian cells, septins exert pivotal functions once the ICB is established and extends^37^, by establishing barriers to impede diffusional spread of cytokinetic proteins^17^, and by scaffolding the assembly of the abscission machinery^37,39^.

To analyze how individual PIPKI isozymes affect septin localization during cytokinesis we assessed the subcellular distribution of SEPT2, a highly abundant paralog present in both hexa- and octamers. In the vast majority of control cells, as well as in cells depleted of PIPKIα or PIPKIβ, SEPT2 was found to be enriched at the ICB (Fig. 1H), with about 5% of total cellular SEPT2 residing on the acetylated tubulin bridge (Fig. 1I). Knockdown of PIPKIγ led to a marked loss of SEPT2 from microtubules at the ICB, which instead remained associated with the plasma membrane of the newly forming daughter cells, at sites adjacent to the bridge membrane (Fig. 1H).

These findings establish PIPKIγ as a novel regulator of anillin and septin organization at the ICB.

### Distinct splice variants of PIPKIγ associate with septins, and colocalize with septin filaments

The carboxyterminal region of human PIPKIγ undergoes alternative splicing, which gives rise to at least five isoforms, termed PIPKIγ-i1-i5 (Fig. 2A). Distinct splice inserts promote PIPKIγ association with select binding partners at specific subcellular localizations^27^, and this mechanism allows for the generation of PI(4,5)P_2_ at specific sites. To identify novel binding partners of these splice variants that might connect to the cytokinetic machinery we performed affinity purification experiments. Surprisingly, glutathione-S-transferase (GST)-fused tails of both PIPKIγ-i3 and -i5, efficiently retained a variety of septin paralogs (Fig. 2B), indicating a specific interaction with septin oligomers. PIPKIγ-i2 and i3 efficiently pulled down the focal adhesion protein talin from mouse brain lysates (Fig. 2B), as expected^31,32^ (Fig. 2A). We then performed alanine scanning mutagenesis along the i3/i5-specific splice insert, and identified two neighboring residues (Y646 and W647) critically involved in septin binding. Accordingly, the tail of a PIPKIγ-i5 mutant (Y646A/Y647A, hereafter referred to as PIPKIγ-i5ΔSB, “deficient in septin binding”) was unable to affinity-purify septins (Fig. 2B). Conclusively, mCherry-PIPKIγ-i5 precipitated myc-SEPT7, as well as SEPT9 or SEPT5 from HEK-293T cell lysates, but not endogenous talin (Fig. 2C, Fig. S2A/B).

**Figure 2:**
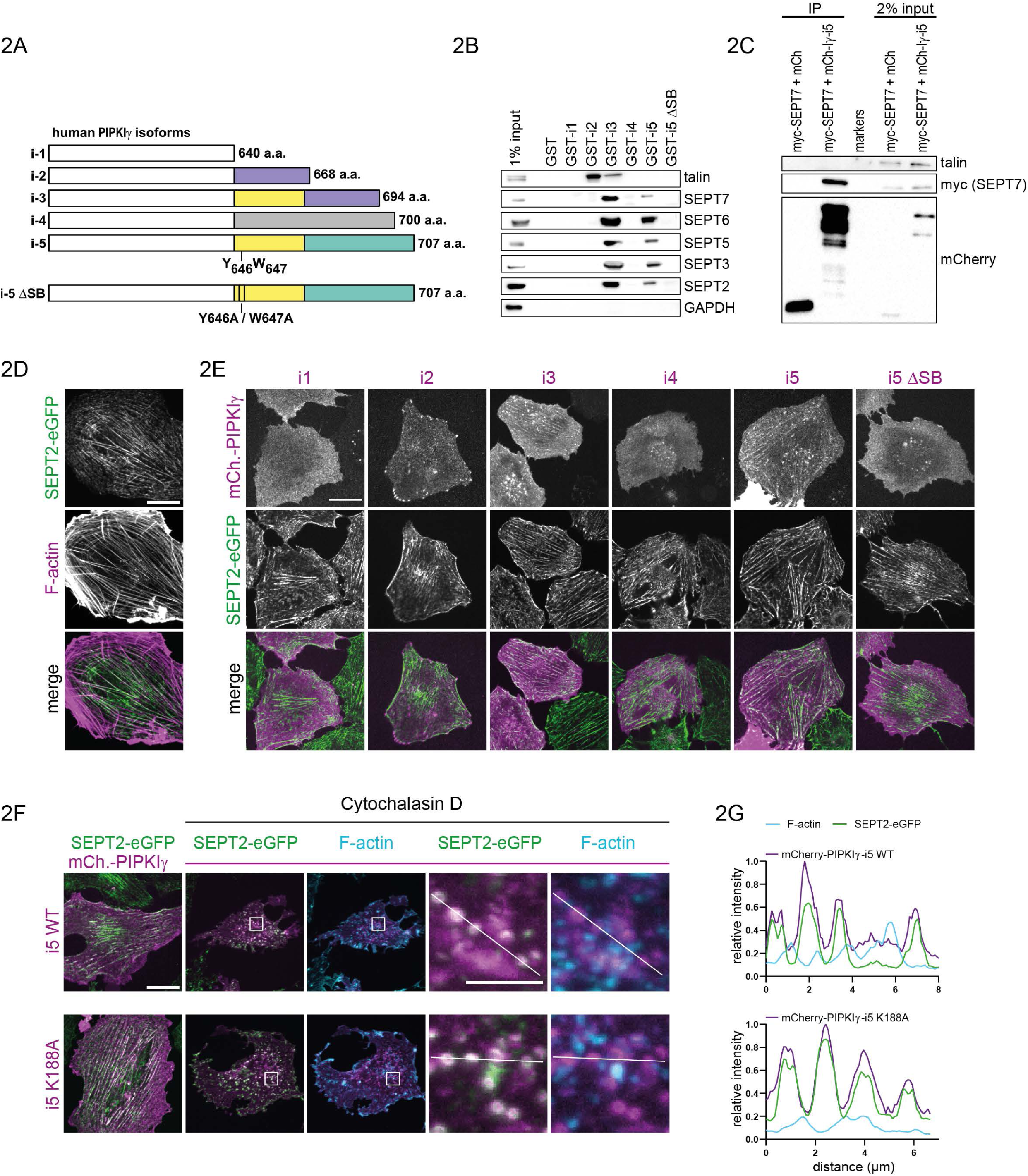
PIPKIγ isoforms i3 and i5 specifically interact with septins. **(A-E)** PIPKIγ isoforms 3 and 5 (i3/i5) interact with septins through two aromatic amino acids (W646 and Y647) harbored in their common splicing insert. **(A)** Schematic representation of C-terminal variations within the tail domains of human PIPKIγ isoforms. A splice insert present in isoforms i2 and i3 (purple) mediates talin binding. Isoforms i3 and i5 share a splice insert (yellow) of unknown function, and analyzed in detail in this study. Note that the kinase core domain common to all isoforms (white box) is not represented in scale. **(B)** Affinity purification experiment on GST-fused kinase tails (aa451 to end) of human PIPKIγ isoforms. Material retained from mouse brain lysates was separated by SDS-PAGE, and analyzed by Western blotting using the indicated antibodies. i5-ΔSB, double mutant (W646A/Y647A) defective in septin binding. **(C)** Co-immunoprecipitation of overexpressed myc-tagged SEPT7 with mCherry (mCh)-tagged PIPKIγ-i5. HEK-293T cells were transiently cotransfected, and lysates were purified on an RFP-affinity resin. The retained material was analyzed by SDS-PAGE and Western blotting using the indicated antibodies. **(D)** Representative confocal image displaying partial overlap between endogenous SEPT2-EGFP and F-actin in NRK49F knock-in cells, stained with phalloidin. Scale bar: 20μm. **(E)** Representative images of live NRK49F knock-in cells expressing SEPT2-EGFP upon transfection of mCherry (mCh.)-tagged PIPKIγ isoforms. Scale bar: 20μm. **(F-G)** PIPKIγ-i5 association with septins is actin-independent, and does not rely on kinase activity. **(F)** Representative confocal images (max intensity z-projection) of NRK49F SEPT2-EGFP knock-in cells transfected with plasmids encoding human mCherry (mCh.)-tagged PIPKIγ-i5, or a kinase-dead mutant (K188A). Cells were incubated with 5 µM cytochalasin D to disrupt actin filaments. Cells were fixed, and F-actin was visualized by AF647-phalloidin. Scale bar: 20 μm, inset: 5 μm. **(G)** Intensity profiles (normalized to the maximum value) of F-actin, SEPT2-EGFP and mCherry PIPKIγ-i5 wild type or K188A, along a line as depicted in (F).

Next, we investigated a potential colocalization of PIPKIγ splice variants with septins. To this end we transfected HeLa cells with HA-tagged versions of PIPKIγ. Notably, when expressed at low levels HA-PIPKIγ-i3 and -i5 were organized in filaments that overlapped with endogenous septin fibers, as visualized by immunostaining of SEPT6 (Fig. S2C). No such colocalization was observed for PIPKIγ- i1 or -i2, or for septin binding-deficient mutant PIPKIγ-i5ΔSB. We corroborated these findings in genome-edited Norwegian Rat Kidney fibroblasts (NRK49F) that express SEPT2-EGFP under its endogenous promoter^40^. In these cells, septins display a characteristic, and prevalent association with actin filaments (Fig. 2D). Again, both mCherry-PIPKIγ-i3 and -i5, but not septin binding-deficient mutant PIPKIγ-i5ΔSB, were found to colocalize with endogenous SEPT2 (Fig. 2E). By contrast, mCherry-PIPKIγ- i1, i2 or i4 displayed no overlap with septins. In line with previous reports i1 was uniformly distributed across the plasma membrane, i2 additionally localized to focal adhesions^31,32^, and i4 was detected in nuclear foci, presumably representing speckles^41^.

F-actin is known to serve as a template for the assembly of septins into filaments^42^, raising the possibility that the colocalization of septins with mCherry-PIPKIγ-i5 might be bridged by actin. We, therefore, treated NRK49F cells with cytochalasin D to disrupt actin filaments. Application of cytochalasin D triggered a rapid collapse of SEPT2-EGFP-containing filaments into rings, as seen before^18^, which retained their colocalization with the kinase, but, importantly, lost their association with F-actin (Fig. 2F/G). Similar observations were made for the catalytically inactive mutant mCherry-PIPKIγ-i5 (K188A).

Together, these findings demonstrate that PIPKIγ splice isoforms i3 and i5 associate with septins, and in cells colocalize with septin filaments or rings, independent of kinase activity, or of the integrity of actin filaments. Given the pivotal role of septins during cell division this suggests that the functions of PIPKIγ during late stages of cytokinesis may be exerted by these splice variants.

### The septin binding splice variants PIPKIγ-i3/i5 organize septins and anillin during cytokinesis

To analyze the subcellular distributions of PIPKIγ-i5, anillin, and septins at different stages of mitosis, we generated a HeLa cell line stably expressing mCherry-tagged PIPKIγ-i5 (Fig. 3A). At anaphase mCherry-PIPKIγ-i5 was homogenously distributed at the plasma membrane. SEPT2 appeared enriched at the equatorial plane, and at the poles of the mother cell, while anillin was found exclusively at the equatorial plane. During transition to telophase mCherry-PIPKIγ-i5 became concentrated at the cleavage furrow, where it aligned with SEPT2 and anillin. Upon completion of furrow ingression mCherry-PIPKIγ-i5 concentrated at the ICB, where it outlined the midbody. Anillin accumulated at the midbody, while SEPT2 started to translocate onto microtubules.

**Figure 3:**
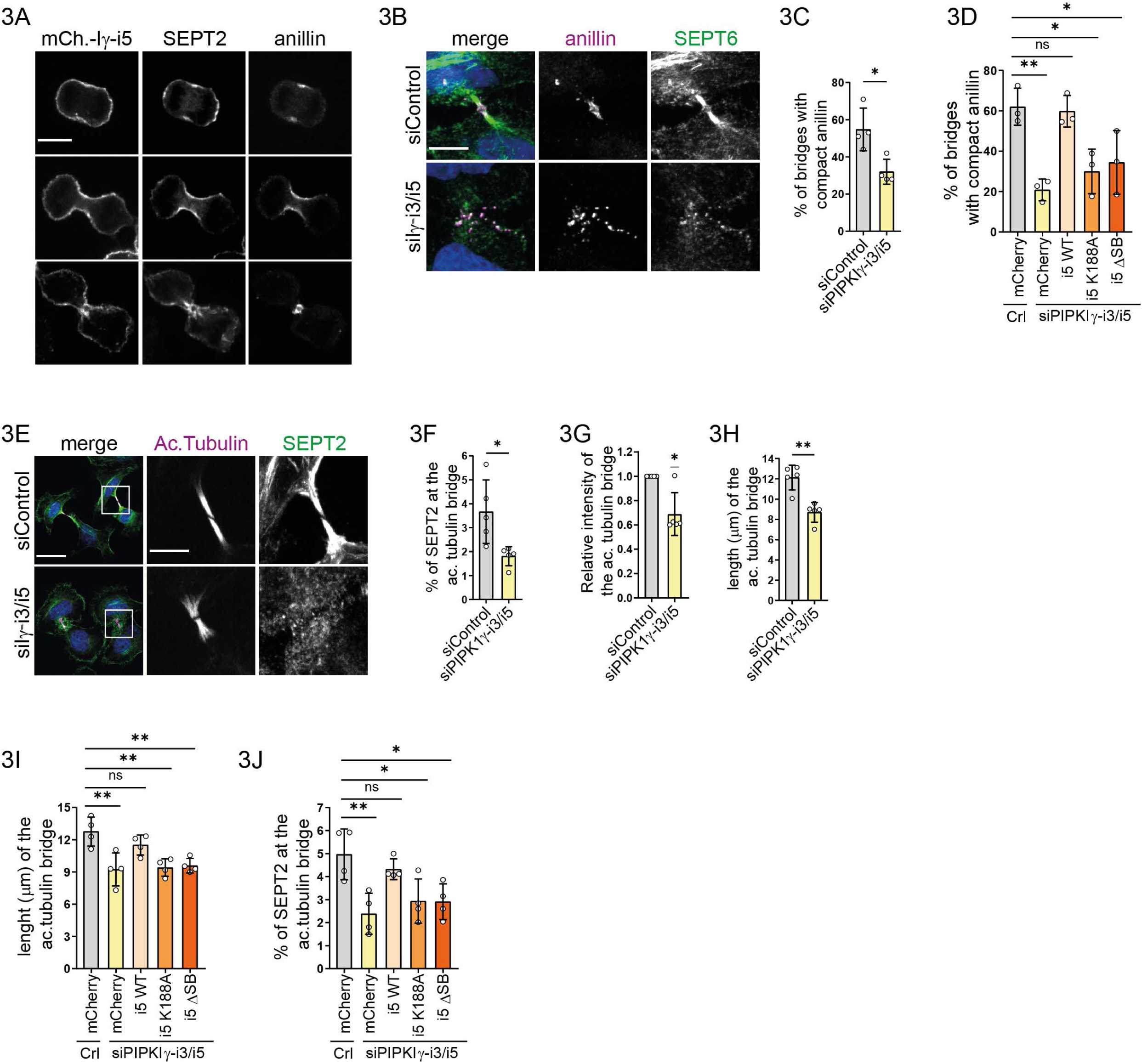
PIPKIγ-i5 localizes to the cleavage furrow, and its regulatory role depends on its kinase activity and its septin binding capability. **(A**) Representative confocal pictures of different mitotic stages of HeLa cells stably expressing mCherry-PIPKIγ-i5, and stained for anillin and SEPT2, scale bar: 10μm. **(B-C)** Selective depletion of PIPKIγ-i3/i5 scatters anillin at the CB. **(B)** Representative confocal pictures (max intensity z-projection) of HeLa cells treated with siRNA against PIPKIγ-i3/i5 or control, synchronized at late cytokinesis, and stained for anillin and SEPT6. Scale bar: 10μm. **(C)** Percentage of ICBs with compact anillin. Data are represented as mean ± SD (n=4, 15-30 bridges were imaged per condition per experiment). Statistics: two-tailed unpaired t test (P= 0.0142). **(D)** PIPKIγ-i5 wild type, but not K188A or ΔSB, rescues anillin accumulation at the midbody. Knock down of PIPKIγ-i3/i5 was performed in HeLa cells stably expressing mCherry or siRNA-resistant mCherry tagged PIPKIγ-i5 wild type, kinase dead (K188A) or ΔSB. The percentage of bridges displaying compact anillin was quantified, and compared to HeLa cells stably expressing mCherry, and treated with control siRNA (see Fig. S3G for representative images). Data are represented as mean ± SD (n=3, 15-30 cytokinetic cells were imaged per condition per experiment). Statistics: One-way ANOVA, followed by Dunnett’s multiple comparison test. Adjusted P values: siControl vs. siPIPKIγ + mCherry, P=0.0023; siControl vs. siPIPKIγ + i5 (WT), P= 0.9962; siControl vs. siPIPKIγ + i5 (K188A), P=0.0121; siControl vs. siPIPKIγ + i5 ΔSB, P=0.0282. **(E-H)** Loss of PIPKIγ-i3/i5 perturbs ICB integrity and impairs SEPT2 recruitment to the ICB. **(E)** Representative confocal images (max intensity z-projection) of HeLa cells treated with siRNA against PIPKIγ-i3/i5 or control, synchronized at late cytokinesis, and stained for acetylated tubulin and SEPT2. Scale bar of merge: 30μm, of grey insets: 10μm. **(F)** Percentage of total SEPT2 detected at the acetylated tubulin bridge, depicted as mean ± SD (n=5, 15-30 cytokinetic cells per condition per experiment). Statistic: two-tailed, unpaired t-test (P=0.0171). **(G)** Relative intensity of the acetylated tubulin bridge. Quantification was performed on average intensity z-projections after background subtraction. Normalized data are represented as mean ± SD (n=5, 15-30 cytokinetic cells per condition per experiment). Statistics: two-tailed, one sample t test (hypothetical value: 1; P: 0.017). **(H)** The length of the acetylated tubulin bridge was measured as max Feret diameter of the ROI delimiting the acetylated tubulin bridge. Data are represented as mean ± SD (n=5, 15-30 cytokinetic cells per condition per experiment). Statistics: two-tailed, unpaired t-test (P=0.0012). **(I)** Expression of PIPKIγ-i5 wild type, but not of kinase dead or ΔSB mutants, rescues the length of the acetylated tubulin bridge (max Feret diameter), mean ± SD (n=4). Statistics: One-way ANOVA, followed by Dunnett’s multiple comparison test. Adjusted P values: siControl vs. siPIPK1γ + mCherry, P=0.0015; siControl vs. siPIPKIγ + i5 (WT), P= 0.3474; siControl vs. siPIPK1γ + i5 (K188A), P=0.0023; siControl vs. siPIPK1γ + i5 (ΔSB), P=0.0036. **(J)** Expression of PIPKIγ-i5 wild type, but not of kinase dead or ΔSB mutants, rescues septin accumulation at the acetylated tubulin bridge (see Figure S3K for representative images). Bar diagram depicts percentage of total SEPT2 at the acetylated tubulin bridge, mean ± SD (n=4, 15-30 cytokinetic cells per condition per experiment). Statistics: One-way ANOVA, followed by Dunnett’s multiple comparison test. Adjusted P values: siControl vs. siPIPKIγ + mCherry, P=0.0026; siControl vs. siPIPKIγ + i5 (WT), P= 0.6814; siControl vs. siPIPK1γ + i5 (K188A), P=0.0156; siControl vs. siPIPK1γ + i5 (ΔSB), P=0.0145.

To investigate whether the disorganization of anillin and septins at the ICB (Fig. 2F-G) could be ascribed to the septin-binding isoforms of PIPKIγ, we selectively depleted PIPKIγ-i3/i5 with an siRNA that targets the common splice insert^33^. This treatment reduced mRNA levels of the two cognate splice variants by about 85%, but did not affect the expression levels of the other three isoforms (Fig. S3A/B). Select depletion of PIPKIγ-i3/i5 increased multinucleation to a similar degree as the general depletion of all PIPKIγ transcripts (Fig. S3C), and was accompanied by a dramatic scattering of anillin from the center of the ICB to its periphery (Fig. 3B/C). To test if this was the consequence of a local loss of PI(4,5)P_2_ at the ICB we artificially increased PI(4,5)P_2_ levels by co-depleting OCRL (also known as Lowe oculocerebrorenal syndrome protein), a 5-phosphatase that is delivered to the ICB by Rab35 prior to abscission to locally erase PI(4,5)P_2_^43^. Knockdown of OCRL alone modestly increased the fraction of cells with compact anillin at their ICB (Fig. S3D-F). Importantly, co-depletion of OCRL together with PIPKIγ-i3/i5 completely rescued the mislocalization of anillin observed upon loss of PIPKIγ-i3/i5 alone (Fig. S3E/F), suggesting that PI(4,5)P_2_ indeed plays a pivotal role for anillin deposition at the midbody. In line with this interpretation anillin dispersion was also rescued by re-expression of an mCherry-tagged, siRNA-resistant wild-type, but not of catalytically inactive mutant PIPKIγ-i5 (Fig. 3D, S3G). Septin binding-defective mutant PIPKIγ-i5ΔSB also failed to rescue anillin dispersion. Failure to rescue anillin dispersion was associated with mistargeting of mutant PIPKIγ-i5: Whereas wildtype mCherry-PIPKIγ-i5 was found enriched at the ICB, in close proximity to endogenous anillin (Fig. 3A, Fig. S3G), mCherry-PIPKIγ-i5 K188A, or ΔSB, were spread along the plasma membrane adjacent to the bridge, similar to dispersed anillin (Fig. S3G). Taken together, these data suggest that septins are required to anchor cognate PIPKIγ variants at the midbody to produce a local pool of PI(4,5)P_2_ that stably anchors anillin.

Given the striking impact of PIPKIγ-i3/i5 on anillin localization we next assessed its role in the dynamic reorganization of the septin cytoskeleton during cytokinesis. To this end we generated a genome-edited cell line expressing EGFP-SEPT6 from its endogenous promotor. We obtained several heterozygous knock-in clones, in all of which endogenous SEPT6 and EGFP-SEPT6 protein levels decreased upon treatment with siRNA (Fig. S4A). Further, EGFP-SEPT6 was successfully incorporated into septin filaments (Fig. S4B), and displayed a high degree of colocalization with endogenous septin paralogs (Fig. S4C). Next, we monitored the subcellular distribution of EGFP-SEPT6 during cytokinesis (Movie 1; representative frames reported in Fig. S3H). Under control conditions EGFP-SEPT6 first accumulated at the cleavage furrow (white asterisk in Fig. S3H). About 30 min after completion of furrow ingression the ICB appeared. At the same time EGFP-SEPT6 started to relocate from the cell cortex onto putative microtubules within the ICB, to remain situated there during bridge elongation until abscission. After abscission the daughter cells displayed perinuclear, sinuous septin fibers that seemed to derive from the ICB (white arrowheads). In cells depleted of PIPKIγ-i3/i5 EGFP-SEPT6 still localized to the cleavage furrow (Movie 2; representative frames reported in Fig. S3H). However, the protein failed to accumulate at the ICB and to translocate onto microtubules, and the resulting daughter cells lacked the prominent septin fibers observed under control conditions. This defect persisted throughout interphase.

Septins are well known to associate with microtubules to regulate their stability and dynamics^16^. Given the dramatic impact of PIPKIγ on septin distribution we investigated the organization of ICB microtubules in more detail. As shown in Fig. 3E depletion of PIPKIγ-I3/i5 induced scattering of septin filaments away from the acetylated tubulin bridge (Fig. 3E/F), similar to observations made upon knockdown of all PIPKIγ isoforms (compare Fig. 1H). The microtubules within the ICB appeared less bundled (Fig. 3E), and recruited less of the microtubule bundling factor PRC1 (Fig. S3I/J). Likewise, microtubule stability was impaired, as assessed by immunostaining against acetylated tubulin (Fig. 3G). Knockdown of PIPKIγ-i3/i5 also significantly reduced the length of the acetylated tubulin bridge (Fig. 3H). Again, wildtype mCherry-PIPKIγ-i5, but neither of the two mutants, could rescue changes in the length of the acetylated tubulin bridge (Fig. 3I), and defects in septin organization (Fig. 3J, Fig. S3K).

In summary, these data demonstrate that distinct splice variants of PIPKIγ are indispensable for the organization of anillin and septins at the ICB, in a process that requires its catalytic activity and its septin-binding capability. The scattering of anillin and septins away from the ICB is accompanied by a loss of microtubule stability, that likely reflects the lack of a local pool of PI(4,5)P_2_.

### Septin-dependent recruitment of PIPKIγ is essential to maintain centralspindlin at the midbody

The tethering of midbody microtubules to the plasma membrane is mediated by centralspindlin, a protein complex composed of MgcRacGAP and mitotic kinesin-like protein 1 (MKLP1)^24^. MgcRacGAP contains an atypical C1-domain with affinity for PI(4,5)P_2_^23^. Based on our observations we hypothesized that the septin-dependent recruitment of PIPKIγ is required for the stable association of centralspindlin with the cell cortex.

To test this hypothesis we assessed the localization of centralspindlin at late stages of cytokinesis. As expected, under control conditions MKLP1 colocalized with the midbody ring, as evidenced by immunostaining together with citron kinase (CIT-K) (Fig. 4A). Notably, knockdown of PIPKIγ-i3/i5 had no impact on the distribution of CIT- K, or on its abundance at the midbody (Fig. 4A/B). Loss of PIPKIγ-i3/i5, thus, does not generally affect midbody composition, or positioning. However, depletion of PIPKIγ-i3/i5 significantly decreased MKLP1 levels at the midbody ring (Fig. 4A/C). This defect was rescued upon expression of wildtype mCherry-PIPKIγ-i5, but not of the catalytically inactive, or of the septin binding-deficient kinase mutant (Fig. 4D, Fig. S5A). Consistent with a primary role of septins in guiding PIPKIγ to the ICB, depletion of SEPT2 impaired the deposition of centralspindlin at the midbody to a similar extent (Fig. 4E/F).

**Figure 4:**
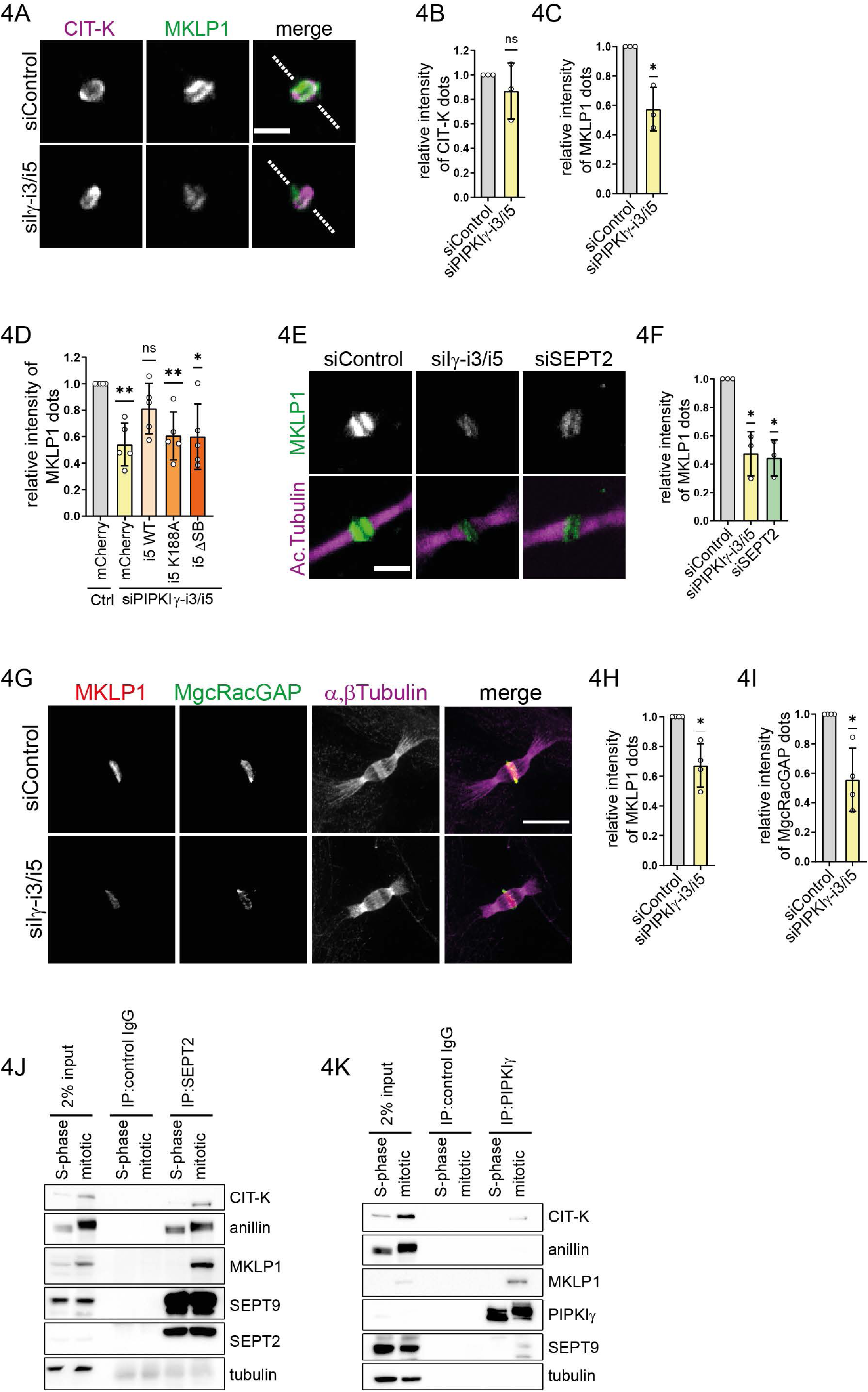
A septin and active PIPKIγ-i5 interacting module regulates the accumulation of centralspindlin at the midbody. **(A-C)** Depletion of PIPKIγ-i3/i5 selectively affect the enrichment of centralspindlin at the midbody. **(A)** Representative confocal images of HeLa cell midbodies upon depletion of PIPKIγ-i3/i5, synchronization at late cytokinesis, and immunostaining of CIT-K and MKLP1. The dashed line indicates the orientation of the cytokinetic bridge. Scale bar: 3 µm. **(B)** Relative intensity of CIT-K **(C)** or MKLP1 dots at the midbody. Quantifications were performed on average intensity z-projections after background subtraction. Normalized data are represented as mean ± SD (n=3, 15-30 cytokinetic cells per condition per experiment). Statistics: two-tailed, one sample t-test (hypothetical value: 1; P values: P=0.4217 in B and P=0.0214 in C). **(D)** Expression of PIPKIγ-i5 wild type, but not of a kinase dead or ΔSB mutants, rescues MKLP1 accumulation at the midbody (see Figure S5A, for representative images). Quantifications were performed on average intensity z-projections after background subtraction. Normalized data are represented as mean ± SD (n=5, 15-30 cytokinetic cells per condition per experiment). Statistics: two-tailed one sample t-test (hypothetical value: 1). P values: siPIPKIγ + mCherry, P= 0.0031; siPIPKIγ + i5 (WT), P= 0.0909; siPIPK1γ + i5 (K188A), P= 0.0081; siPIPK1γ + i5 (ΔSB), P= 0.0224. **(E-F)** Depletion of SEPT2 phenocopies the loss of MKLP1 upon depletion of PIPK1γ-i3/i5. **(E)** Representative confocal images of HeLa cell midbodies upon knockdown of PIPKIγ-i3/i5 or SEPT2, synchronization at late cytokinesis and immunostaining of MKLP1 and acetylated tubulin. Scale bar: 3µm. **(F)** Relative intensity of MKLP1 dots at the midbody. Quantification was performed on average intensity z-projections after background subtraction. Normalized data are represented as mean ± SD (n=3; 15-30 cytokinetic cells were imaged per condition per experiment). Statistics: two-tailed one sample t-test (hypothetical value: 1). P values: siPIPKIγ-i3/i5 P=0.0279; siSEPT2 P=0.0167. **(G-I)** Ultrastructure expansion microscopy (U-ExM) images indicating defects in centralspindlin accumulation at the midbody upon depletion of PIPKIγ-i3-i5. **(G)** Cells were synchronized at late cytokinesis and expanded using the U-ExM method (Gambarotto et al., 2019). Expanded gels were immunostained for MKLP1, MgcRacGAP, and α-/β-tubulin, and imaged on a spinning disk confocal microscope. Representative pictures (max intensity projections of 21 slices with 1 µm spacing) are shown. Scale bar: 5μm. **(H)** Relative intensity of MKLP1 and **(I)** MgcRacGAP at the midbody. Quantifications were performed on average intensity z-projections after background subtraction. Data are represented as mean ± SD (n=4; 20-25 bridges were imaged per condition per experiment. Statistics: two-tailed one sample t-test (hypothetical value: 1; P values: P=0.0203 in H and P =0.0256 in I). **(J-K)** Septins and PIPKIγ interact with components of the midbody. **(J)** SEPT2 or **(K)** PIPKIγ were immunoprecipitated from lysates of synchronized HeLa cells. The affinity-purified material was separated by SDS-PAGE and analyzed by Western blotting using the indicated antibodies.

Next, we aimed at gaining insight into the nanoscale organization of septins, microtubules and centralspindlin at the midbody. The detection of midbody proteins by conventional indirect immunofluorescence microscopy is complicated by the dense packing of individual components in this organelle, which allows only limited access for antibodies. To overcome this problem, we used a variation of expansion microscopy namely ultrastructure expansion microscopy (U-ExM)^44^, for which the specimen is physically expanded prior to antibody labeling^45^. At cytokinesis control cells exhibited bundles of antiparallel microtubules along the ICB that overlapped at the midbody (Fig. S5B). Septins were not found at the midbody itself, but formed two rings that aligned with two secondary ingression sites at its flanks (Fig. S5B). Additionally, they colocalized with microtubules that protruded away from the secondary ingression sites, into the developing daughter cells. Upon depletion of PIPKIγ-i3/i5 septins underwent a dramatic redistribution. They were mostly dispersed from the ICB, and only sparsely aligned with microtubules. Instead of being organized in continuous, straight fibers they formed short rods and curly filaments in proximity to the ICB at the periphery of the daughter cells. Next, we assessed the distribution of centralspindlin by U-ExM. In control cells centralspindlin localized to a confined region at the midbody, where microtubules overlapped. In PIPKIγ-i3/i5-depleted cells total centralspindlin levels at this locale were significantly reduced, akin to our findings in non-expanded specimen (Fig. 4G-I).

The PI(4,5)P_2_-dependent recruitment of effector proteins to the plasma membrane is frequently fostered by complex formation between the effector protein itself, and PI(4,5)P_2_-synthesizing enzymes^35^. This raises the possibility that centralspindlin might interact with PIPKIγ. We, thus, screened for binding partners of septins and PIPKIγ in cytokinetic HeLa cells. We identified the centralspindlin component MKLP1 in immunoprecipitates of SEPT2 and of PIPKIγ (Fig. 4J/K). Citron kinase, a known interactor of centralspindlin^46^, was detected in both precipitates as well. Anillin, however, efficiently copurified only with septins, but not with PIPKIγ. This suggests that the association of septins with PIPKIγ is not bridged by anillin.

Given the pivotal function of PIPKIγ-i3/i5 on midbody organization we next investigated its impact on the distribution of PI(4,5)P_2_ in synchronized cells. In control cells PI(4,5)P_2_ was found at the outlines of the ICB, occasionally enriched at the MgcRacGAP-positive midbody (Fig. S5C). Knockdown of PIPKIγ-i3/i5 greatly reduced the levels of MgcRacGAP at the ICB, as expected. At the same time PI(4,5)P_2_ appeared more diffusely distributed across a broader area, and was enriched in filopodia extending between the daughter cells in parallel to the bridge. Line scans perpendicular to the axis defined by the ICB revealed two major peaks of PI(4,5)P_2_ intensity, adjacent to the central midbody (Fig. S5C). Loss of PIPKIγ-i3/i5 did not have a major impact on peak intensities (Fig. S5D), indicating that in absence of PIPKIγ-i3/i5 PI(4,5)P_2_ homeostasis at the ICB is not globally affected. However, under knockdown conditions the peaks were broader, and were further separated as compared to controls. In line with the loss of centralspindlin from the midbody (Fig. 4A/E/G) this may reflect a widening of the ICB^23^.

Taken together, these data data reveal that septin-dependent recruitment of PIPKIγ-i3/i5 is essential to maintain centralspindlin at the midbody and this is essential for the successful completion of cytokinesis.

## Discussion

Almost two decades ago PI(4,5)P_2_ was identified as a key factor required for cytokinesis ^10^, and over time a steadily increasing number of components of the cytokinetic machinery has been identified that depend on this phosphoinositide for their recruitment to the plasma membrane^9^. Yet, the molecular mechanisms underlying its synthesis at the cleavage furrow and at the maturing ICB have remained elusive. Our study provides first and important insight into how PIPKIγ splice variants are recruited to the midbody, to generate a pool of PI(4,5)P_2_ dedicated to maintain centralspindlin (Figure 5). We demonstrate that a distinct splice insert promotes the association of PIPKIγ with septins in interphase, as well as in mitotic cells. In line with a function during cell division, depletion of PIPKIγ causes multinucleation to a similar extent as loss of septins^17,47^, and leads to a dramatic scattering of anillin and septins away from the ICB during cytokinesis. Further, we identify centralspindlin as a common binding partner of septins and PIPKIγ, and show that loss of either, septins or PIPKIγ, impairs centralspindlin accumulation at the midbody.

**Figure 5:**
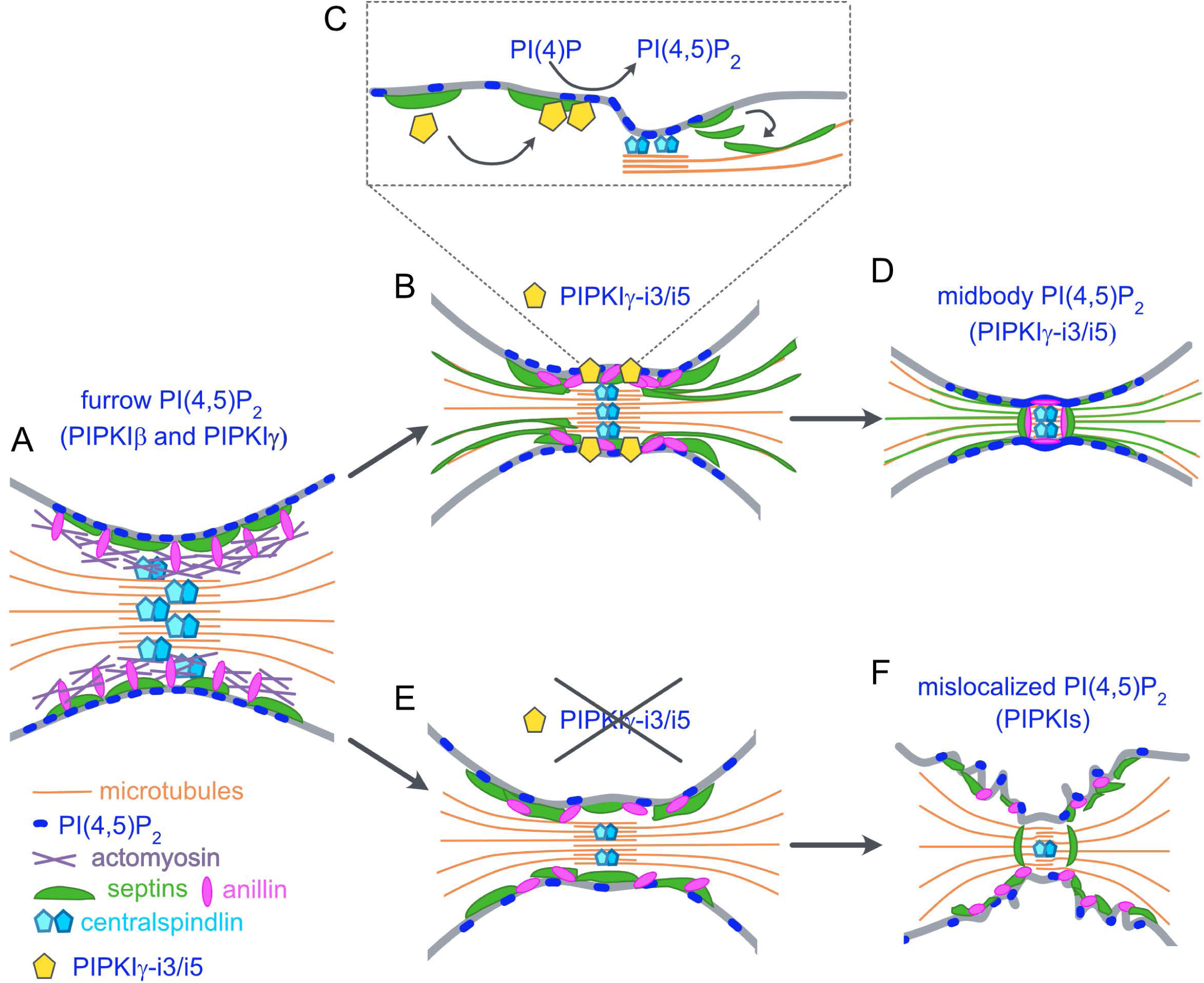
Model summarizing the role of PIPKIγ-i3/i5 during cytokinesis. **(A)** Anillin (pink) and septins (green) are enriched at the cleavage furrow, where they act as scaffolds to initiate and sustain actomyosin-mediated constriction. **(B)** Through interaction with septins, PIPKIγ-i3/i5 (yellow pentagons) are recruited to the furrow, where they generate a local pool of PI(4,5)P_2_. This pool is required for the maintenance of anillin and centralspindlin (turquoise) at the nascent midbody. **(C)** The centralspindlin-mediated tethering of the furrow membrane to microtubules possibly facilitates the translocation of septins onto bridge microtubules. **(D)** Presence of septins on bridge microtubules favors their bundling, while the ICB matures. **(E)** Depletion of PIPKIγ-i3/i5 disrupts the localized synthesis of PI(4,5)P_2_ at the nascent midbody, impairing centralspindlin tethering to the plasma membrane. **(F)** Mislocalized synthesis of PI(4,5)P_2_ may contribute to the scattering of anillin and septins. Disorganized septins (and anillin) further affect the maintenance of centralspindlin at the midbody, and the maturation of the ICB.

Based on our observations we propose the following model: In cytokinetic cells anillin-associated septins recruit PIPKIγ-i3/i5 to the constricting cleavage furrow. Already at that stage PIPKIγ, likely together with PIPKIβ^14^, contributes to cytokinetic progression by fuelling PI(4,5)P_2_ synthesis at the furrow (Figure 5A). As soon as the cleavage furrow encounters spindle microtubules at the ICB (Figure 5B), septin-bound PIPKIγ-i3/i5 generate an additional, local pool of PI(4,5)P_2_. This pool is essential to retain septins, anillin and centralspindlin at the nascent midbody (Figure 5C). Centralspindlin-dependent tethering of microtubules to the furrow membrane likely facilitates the translocation of septins onto microtubules, where septins support microtubule stability, and favor their bundling during ICB elongation (Figure 5D). In absence of PIPKIγ-i3/i5 anillin and septins are scattered away from the ICB, and centralspindlin anchorage at the midbody is impaired (Figure 5E). Mislocalized synthesis of PI(4,5)P_2_ may facilitate clustering of anillin and septins at sites apart from the ICB. Microtubules are less stablilized and less bundled, and the bridge fails to elongate (Figure 5F).

According to this model we suggest that the assembly of septins, PIPKIγ and centralspindlin establishes an important checkpoint that arrests cytokinesis progression until the ingressed cleavage furrow has successfully encountered the midbody. Conclusively, depletion of septins^17,18^, manipulation of PI(4,5)P_2_ levels^10^, expression of anillin mutants deficient in septin binding^12,37^, or of membrane binding-defective MgcRacGAP C1 domain mutants^23^ result in similar defects, i.e. furrow instability, eventual retraction of the cleavage furrow, and ultimately multinucleation.

Based on rescue experiments we conclude that this late function of PIPKIγ strictly requires kinase activity. However, we were unable to visualize significant changes in PI(4,5)P_2_ levels at the ICB upon kinase depletion. Under control conditions we detected enrichment of PI(4,5)P_2_ at the midbody itself only occasionally. This might indicate that the PIPKIγ-dependent pool of PI(4,5)P_2_ accumulates only very transiently, or locally. Alternatively, its detection might be hindered due to the tight association and/or local enrichment of lipid effectors, which instantly mask newly synthesized PI(4,5)P_2_. In support of the latter conclusion similar complications arise during the detection of other PI(4,5)P_2_ pools, such as those generated at focal adhesions^31,32^, on sorting endosomes^33^, or at sites of clathrin-mediated endocytosis^48^.

Of note, we observed significant changes in the distribution of PI(4,5)P_2_ at the ICB. In control, as well as in PIPKIγ-i3/i5-depleted cells PI(4,5)P_2_ outlined the boundaries of the ICB, resulting in two major peaks detected in vicinity of the midbody. Upon loss of PIPKIγ the PI(4,5)P_2_ staining appeared more diffuse, and the distance between the two peaks increased, suggesting a widening of the ICB, which likely is caused by the loss of centralspindlin^23^.

Consistent with defects in the nanoscale distribution of PI(4,5)P_2_ in kinase-depleted cells we observed massive scattering of anillin and septins away from the ICB. As anillin requires septin binding for its concentration at the ICB^37^ its loss could be an indirect effect of septin dispersal. Indeed, displaced anillin and septins remain colocalized in regions peripheral to the ICB. Furthermore, this might be a consequence of decreased local PI(4,5)P_2_ levels, as co-depletion of OCRL, a 5-phosphatase involved in late-stage clearance of PI(4,5)P_2_ from the ICB^43^, reverts the loss of anillin from the midbody. Previous reports based on super-resolution imaging suggested that septins and anillin form ring-like arrays along the ICB to promote its elongation ^37^. In support, we find that in kinase-depleted cells, i.e. when anillin and septins are scattered away from the ICB, the acetylated tubulin bridge is significantly shorter. As this defect can only be rescued by wildtype, but not by septin binding-deficient, or catalytically-inactive variants of PIPKIγ, our data implicate septin-dependent synthesis of PI(4,5)P_2_ by PIPKIγ also in ICB maturation.

To gain insight into the nanoscale organization of microtubules at the ICB we employed super-resolution expansion microscopy. We demonstrate here for the first time that U-ExM is a powerful technique ideally suited to resolve individual structures in crowded environments, such as the midbody. Our analyses revealed that microtubules interdigitate at the midbody in wild-type, as well as in kinase knockdown cells, and independent of local centralspindlin levels. However, depletion of PIPKIγ completely displaced septins from ICB microtubules in regions distal to the midbody. Consistent with the established roles of septins in regulating microtubule organization^16^, kinase-depleted cells displayed significantly reduced levels of acetylated tubulin and PRC1 at the bridge, indicating defects in microtubule stability and bundling, respectively. Future studies will need to address the underlying molecular mechanisms in more detail.

To visualize the dynamic redistribution of septins during mitosis we monitored eGFP-tagged SEPT6 in genome-edited cells by live-cell imaging. In control cells septins translocate from the cell cortex onto bridge microtubules, as identified by colabeling with acetylated tubulin in fixed specimen, as soon as furrow ingression is completed. They then remain associated with these microtubules as sinuous fibers, even after separation of the daughter cells. Depletion of PIPKIγ-i3/i5 does not impair the association of eGFP-SEPT6 with the ingressing cleavage furrow, but completely abrogates septin association with microtubules. This defect persists even after abscission, indicating that the underlying mechanism not only acts during mitosis, but also in interphase, or that the association of microtubules with septins is a property inherited by the daughter cells.

Midbody microtubules are tethered to the plasma membrane by the MgcRacGAP subunit of centralspindlin, which bears an atypical C1 domain with specificity for PI(4,5)P_2_^23^. Our findings suggest that centralspindlin anchorage at the midbody is mediated by a tight functional interplay between septins and PI(4,5)P_2_, as synthesized by septin-associated PIPKIγ. Accordingly, we find that depletion of septins, or of PIPKIγ impede centralspindlin deposition at the midbody to a similar extent. MgcRacGAP’s C1 domain is dispensable for the localization of centralspindlin *per se*, as shown for variants lacking the C1 domain, or for mutants deficient in PI(4,5)P_2_-binding^23^. We conclude that its anchorage at the midbody also relies on a septin scaffold that needs to assemble at the ICB. As even by expansion microscopy we were unable to detect septins at the midbody itself, we conclude that septins rather assist centralspindlin concentration by acting as an additional anchor, and/or diffusion barrier to prevent its premature loss from the midbody. At the same time, such a diffusion barrier might aid the concentration of PI(4,5)P_2_ along the ICB^49^.

The motor subunit of centralspindlin, MKLP1, has been shown to associate directly with Arf6 to promote its recruitment to the midbody^50^. Arf6 is a small GTPase required for the completion of cytokinesis^51^, and known to directly activate PIPKIγ^52^. This, thus, raises the intriguing possibility that the septin/PIPKIγ-dependent centralspindlin deposition at the midbody acts as a feed-forward mechanism to locally enhance PI(4,5)P_2_ production, and to thereby foster recruitment of further midbody components, such as exocyst^53,54^. One of the key functions of centralspindlin throughout cytokinesis is the regulation of Rho GTPases, which is exerted through MgcRacGAP^24^. Loss of centralspindlin from the midbody in PIPKIγ-depleted cells might, thus, lead to a mis-regulation of GTPase activities at the midbody and along the ICB. Consistently, RhoA hyperactivity has been observed in PIPKIγ knockout cells^55^. Hyperactive RhoA, in turn, is known to stimulate the activities of type I PIPKs^56^. This mechanism could, thus, induce PI(4,5)P_2_ production by other isoforms at aberrant sites, and might explain why loss of PIPKIγ does not induce overt alterations in the overall PI(4,5)P_2_ content of the ICB.

## Acknowledgements

We thank Heike Stephanowitz and Fan Liu (FMP Berlin) for mass-spectrometric analyses.

## Author contributions

G.R., S.R., N.J. and N.H. carried out the experiments, C.S., M.L. and H.E. helped with image analysis, F.H. determined mRNA levels upon knockdowns of PIPKIγ isoforms. M.K. designed the project and wrote the manuscript.

## CDeclaration of interests

The authors declare no competing interests.

## Movies

**Movie 1: Dynamic rearrangement of EGFP-SEPT6 during cell division in control cells.**

HeLa EGFP-SEPT6 knock-in were treated with control siRNA, synchronized, and imaged throughout cytokinesis with a confocal microscope. Imaging was started 7,5h after thymidine release, and carried out with a frame rate of 10 minutes. Time stamp: hh:mm; scale bar, 15µm.

**Movie 2: Dynamic rearrangement of EGFP-SEPT6 during cell division in PIPKIγ-i3/i5-depleted cells.**

HeLa EGFP-SEPT6 knock-in were treated with siRNA targeting PIPKIγ-i3/i5, synchronized, and imaged throughout cytokinesis with confocal microscope. Imaging was started 7,5h after thymidine release, and carried out with a frame rate of 10 minutes. Time stamp: hh:mm; scale bar, 15µm.

## Material and Methods

### Plasmids and siRNAs

For mammalian expression constructs the coding sequences (CDSs) of human PIPKIγ isoforms 1 to 5 (PIPKIγ-i1-i5) (Xu et al., 2014), and of i5 mutants were inserted into pcDNA3.1(+)-based vectors, resulting in the expression of N-terminally mCherry- or HA-tagged proteins. The mutant PIPKIγ-i5 K188A carries the previously described mutation within the kinase core domain, which renders the kinase inactive (Krauss et al., 2006). The mutant deficient in septin binding, PIPKIγ-i5 ΔSB, was generated by mutating Y646 and W647 of human PIPKIγ-i5 into alanine by site-directed mutagenesis (Y646A/W647A). SiRNA-resistant PIPKIγ-i5 and -i5 mutants were created by introducing four silent mutations within the sequence targeted by the siRNA against PIPKIγ-i3/i5 as follows: 5’-<u>C</u>GA<u>C</u>GG<u>C</u>AG<u>A</u>TACTGGATT-3’. For viral constructs the resulting CDSs were subsequently inserted into pLIB-CMV-mCherry-IRES-Puro. For bacterial expression constructs the CDSs of the tail domains of PIPKIγ-i1-i5 and -i5 ΔSB (aa 451 to end) were inserted into pGEX-4T1, allowing for the expression of N-terminally tagged glutathione-S-transferase (GST) fusion proteins.

All siRNAs used in this study were purchased from Sigma-Aldrich, and had 3’-dTdT overhangs. For silencing the following siRNAs were used, targeting the human sequences: OCRL 5’-GAAAGGAUCAGUGUCGAUA-3’^43^, SEPT2 5’-GCCCUUAGAUGUGGCGUUU-3’^57^, siSEPT6 5’-CCUGAAGUCUCUGGACCUAGU-3’^17^, siPIPKIγ 5’-GAGGAUCUGCAGCAGAUUA-3’, siPIPKIγ-i3/i5 5’-GGAUGGGAGGUACUGGAUU-3’^33^. On-Target Plus siRNA smart pools (Dharmacon) were used to silence PIPKIα (L-004780-00-0010) or PIPKIβ (L-004058-00-0010). The control siRNA used throughout the study corresponded to the scrambled µ2-adaptin sequence 5’-GUAACUGUCGGCUCGUGGU-3’.

### Generation of antibodies

An antibody specifically recognizing PIPKIγ was raised in rabbits immunized with recombinant His-tagged PIPKIγ (aa451-668; Krauss et al., 2006). The same recombinant protein was covalently coupled to CNBr-coupled sepharose and used for affinity purification. An antibody specifically detecting SEPT6 was raised in rabbits immunized with a peptide comprising aa413-327 of SEPT6 (CAGGSQTLKRDKEKKN), and affinity-purified on the same epitope.

### Cell lines

HeLa and HEK-293T cells were obtained from American Type Culture Collection (ATCC), and not used beyond passage 30 from original derivation. The genome-edited NRK49F SEPT2-eGFP knock-in cell line has been described previously^40^. HeLa cells were cultured in Dulbeccós modified Eaglés medium (DMEM) containing 1g/L D-glucose and phenol red (PAN Biotech), supplemented with 10% (vol/vol) heat-inactivated fetal bovine serum (FBS, Gibco), 2mM L-glutamine (Gibco), 50μg/mL penicillin-streptomycin (pen-strep, Gibco). Stably transfected HeLa cells were generated by viral transduction, and maintained under selection pressure by additionally supplementing the above described medium with 1 µg/mL puromycin (Invivogen). HEK-293T were cultured in DMEM containing 4,5g/L D-glucose, phenol red, L-glutamin (Gibco), supplemented with 10% FBS and 50μg/mL pen-strep. NRK49F SEPT2-eGFP cells were cultured in DMEM containing 4,5g/L D-glucose (Gibco), supplemented with 10% FBS and 2mM L-glutamine. All cell lines were cultured at 37 °C and 5% of CO_2_, and regularly tested for mycoplasma contaminations.

### Generation of a EGFP-SEPT6 knock-in cell line

Endogenous tagging of the SEPT6 N-terminus with EGFP was achieved via the CRISPR/Cas9 technology^58^. In brief, the primers 5’-CACCGCATCGCTCCTGCGTCCGCCA-3’ and 5’-AAACTGGCGGACGCAGGAGCGATGC-3’ were annealed and subcloned into px458-pSpCas9(BB)-2A-GFP (Addgene) using the BpiI restriction site to generate a guide RNA. In a second donor vector the expression cassette was exchanged with the CDS of EGFP inserted between two homology regions (HR) consisting of original genomic sequences ∼ 1000 bp upstream of the SEPT6 ATG (5’HR) and ∼1000 bp downstream of the SEPT6 ATG (3’HR). The stop codon of EGFP was exchanged with two codons encoding for a Gly-Ser linker. Design and cloning of the donor vector was performed with the NEBuilder assembly tool and NEBuilder HiFi DNA assembly cloning kit, respectively, according to the manufacturer instructions. HeLa cells were co-transfected with px458-pSpCas9(BB)-2A-GFP containing the guide sequence, and with donor vector. 72h later eGFP-expressing HeLa were sorted into 96-well plates at a density of one cell per well, using a fluorescence-activated single cell sorter (BD FACSAria). Growing colonies were expanded, and tested for the expression of eGFP-SEPT6 by automated live cell imaging and Western blotting using anti-SEPT6 and anti-GFP antibodies. The expression of eGFP-SEPT6 in selected clones was further validated by siRNA-mediated depletion of SEPT6, by immunoprecipitation of GFP, and by immunocytochemistry.

### siRNA-mediated gene silencing

To silence PIPKIα, PIPKIβ, PIPKIγ and PIPKIγ-i3/i5, two rounds of ∼48h knock-down each were performed with 50 nM siRNA, using JetPRIME (Polyplus) as transfection reagent according to the manufactureŕs instructions. For immunocytochemistry (ICC) and Ultrastructure expansion microscopy (U-ExM, see below) cells were seeded on matrigel (Corning)-coated glass coverslips in a 12-well plate for the second round of knock-down. For live-cell imaging of eGFP-SEPT6, the second round of knock-down was performed in matrigel-coated 8 well glass-bottom slides (ibidi). Cells were analyzed ∼48h after the second round of transfection. Silencing of SEPT2 and SEPT6 was achieved with one round of knock-down for ∼48h with 100 nM siRNA and JetPRIME as transfection reagent according to the manufactureŕs instructions.

For the co-depletion of PIPKIγ-i3/i5 and OCRL, two rounds of knock-down were performed, with 50 nM siPIPKIγ-i3/i5 + 50 nM siOCRL. In the same experiment, the single depletions of PIPKIγ-i3/i5 and OCRL were performed with 50nM targeting siRNA + 50nM siControl, while control cells were treated with 100 nM siControl at each round.

### Plasmid overexpression

Transfection of HeLa was performed with JetPRIME (Polyplus), according to manufactureŕs instructions. The medium was exchanged after 6 hours of transfection, and the cells were processed after 24h of transfection. For the overexpression of PIPKIγ isoforms, cells were seeded on matrigel-coated glass coverslips in a 12-well plate, and transfected with 0.5 μg of DNA per well.

Transfection of NRK49F SEPT2-eGFP was performed with Lipofectamine 3000 (Thermo-Fisher), following the manufactureŕs instructions. Cells were seeded on glass coverslips in a 6-well plate, transfected on the following day, and used for experiments 48h post transfection. For immunoprecipitation assays, or for virus production, HEK-293T cells were transfected using calcium phosphate. In brief, DNA was mixed with 0.12 M CaCl_2_ in 0,1xTE buffer (1 mM TRIS, 0.1 mM EDTA pH 8.0) and incubated for 5 min at room temperature. The same volume of 2X HBS (280 mM NaCl, 10 mM KCl, 1.5 mM Na2HPO4, 12 mM dextrose, 50 mM HEPES pH 7.0) was added while stirring. After 20 min of incubation at room temperature, the DNA solution (1mL for a 10 cm dish) was added to the cells.

### Cytochalasin D treatment and phalloidin staining in NRK49F SEPT2-eGFP

Cells were incubated for 30 min in full medium containing 5 µM cytochalasin D. After a quick rinse with PHEM buffer (60 mM PIPES, 25 mM HEPES, 2 mM MgCl_2_, 10 mM EGTA, pH 6.9), cells were fixed with 4 % PFA in PHEM buffer at 37°C for 15 min. Excess of PFA was quenched with 50 mM NH_4_Cl in PHEM buffer for 10 min on a shaking platform. Cells were washed three times for 5 min on a shaking platform, and stored for imaging, or processed for phalloidin staining.

For phalloidin staining, cells were permeabilized with 0.3 % Triton-X 100 in PHEM buffer for 5 min. Next, samples were blocked with 4 % horse serum and 1 % BSA in PHEM buffer for one hour. Coverslips were incubated with 1 µM AlexaFlour-647-coupled phalloidin (Thermo-Fisher) in 1 % BSA / PHEM buffer for 30 min, and then washed three times for 10 min in PHEM buffer on a shaking platform. Coverslips were mounted in FluoromountG (Invitrogen).

### Immunocytochemistry

HeLa cells seeded on matrigel-coated glass coverslips were fixed with 4% PFA/ 4% sucrose, or with 2% PFA/ 2% sucrose (for septin stainings) in PBS for 15 minutes at RT, and subsequently washed three times with PBS. Cells were permeabilized with washing buffer (20 mM Hepes pH 7,2, 150 mM NaCl, 0.1% Triton X-100) for 15 minutes, and then blocked with goat serum dilution buffer (GSDB: 20mM Hepes pH7.2, 150 mM NaCl, 0.1% Triton X-100, 10 % goat serum) for 20 minutes. Incubation with primary antibodies, diluted in GSDB, was carried out at RT for 1h, and the excess of antibody was removed by three washes of 10 minutes each with washing buffer. Cells were subsequently incubated with AlexaFluor-conjugated secondary antibodies diluted in GSDB for 1h at RT. After three washes in washing buffer coverslips were incubated for 5 minutes with 1 µg/mL of DAPI in PBS, and mounted on microscope glass slides with Immu-Mount (Thermo-Fisher). For the staining of anillin, goat serum was replaced by donkey serum throughout the protocol.

Plasmalemmal PI(4,5)P_2_ was visualized using purified, recombinant pleckstrin homology (PH) domain of PLC-δ1. Cells were fixed with 2% PFA, 2% sucrose + 1% glutaraldehyde (GA) in PBS for 20 minutes at RT. Excess of fixative was quenched by three washes with 50 mM NH_4_Cl in PBS. Permeabilization and blocking were performed concomitantly with a first round of incubation with 0.25 μg/mL of His_6_-tagged EGFP-PH-PLCδ1 in 0.5% saponin, 1% BSA in PBS for 30 minutes at RT. Subsequently, cells were incubated (without washing in between) again with 0.25 μg/mL of His_6_-tagged EGFP-PH-PLCδ1 diluted in 1% BSA in PBS for 30 min at RT. Subsequently, coverslips were washed three times with PBS, and incubated with primary antibody (rabbit-anti-GFP, Abcam) in 1% BSA, 10% GS in PBS for 1h at RT. Excess of antibody was removed by three washes with PBS. The secondary antibody (AlexaFluor488-coupled goat-anti-rabbit) was diluted in 1% BSA, 10% GS in PBS and incubated at RT for 30 minutes. Excess of antibody was removed by four washes with PBS, supplemented with 1 µg/mL of DAPI for the final wash before mounting. To visualize PI(4,5)P_2_ at the ICB by antibody labeling (mouse-anti-PI(4,5)P_2_, Echelon-Z), fixed cells were permeabilized in 0,5% saponin, 1%BSA in PBS for 30 minutes at RT. Coverslips were incubated with primary antibodies diluted in 1% BSA, 10% GS in PBS for 2h at RT. Excess of antibodies was removed by three washes in PBS, and coverslips were incubated with AlexaFluor-coupled secondary antibodies diluted in 1% BSA, 10% GS in PBS for 1h at RT. Coverslips were subsequently washed three times with PBS, once with PBS supplemented with 1 µg/mL of DAPI, and finally mounted.

### Cell synchronization

For microscopy-based assays, 2 mM thymidine was applied in full medium to stall the cells at S-phase. 24h later thymidine was removed by four washes of one minute with PBS, and cells were incubated for 7,5h in fresh medium. Then, fresh medium supplemented with 20ng/mL of nocodazole was added. After 4h cells were carefully washed four times for one minute with full medium pre-warmed at 37°C, and subsequently allowed to proceed to cytokinesis by incubation for 1.5h in full medium, before fixation. For live cell imaging the nocodazole block was omitted.

For immunoprecipitation, the nocodazole block was applied overnight by supplementing the medium with 40 ng/mL of nocodazole. The following day cells were collected by mitotic shake-off, and washed four times in full medium by centrifugation at 300 x *g* for 5 min. Cell were then re-plated in full medium, and allowed to proceed to cytokinesis for 90 min. Cells were collected by gentle shake-off, and harvested by centrifugation at 300 x *g* for 5 min. Before lysis, cells were washed once in PBS. Cells stalled at S-phase for 48h served as controls, and were harvested by trypsinization.

### Microscopy and image analysis

Immunostained HeLa cells were routinely imaged with a Zeiss confocal spinning disk microscope (Yokogawa CSU22, Hamamatsu EMCCD camera), using a 60x immersion oil objective (1.4 NA). At least 12 images were acquired per condition, per experiment. For each image of cytokinetic cells, a stack of 21 pictures within the z-plane (z-stack), with a spacing of 0.2μm was acquired. Cells stained with purified His_6_-tagged EGFP-PH-PLCδ1 domain (fig. S1E), and knock-in EGFP-SEPT6 cells (fig. S4C) were imaged within a single z-plane by focusing on the plasma membrane, or on prominent septin filaments, respectively. For the experiment depicted in Fig. 1A-E and Fig. S1B, semi-automated epi-fluorescent imaging was conducted with a Nikon Eclipse Ti microscope (illumination: CoolLED, pE4000, prime95B sCMOS camera) operated by NIS-Elements software, using a 20x air objective (0.75 NA). Tile scans of 1.3 mm^2^ were produced by stitching together images automatically acquired around a chosen point. Four tile scans were acquired per condition, per experiment. Epi-fluorescent pictures of HeLa cells transfected with PIPKIγ isoforms (Fig. S2C) were acquired with the same microscope, using a 40x immersion oil objective (1.3 NA). Live cell imaging of eGFP-SEPT6 cells was performed on a spinning disk Nikon Eclipse Ti microscope (Yokogawa CSU-X1 and EMCCD Camera), operated by NIS-Elements software, with a 40x air objective (0. 95 NA). Imaging was initiated 7,5h after thymidine release, and was carried out overnight, with a frame rate of 10 minutes. Cells were kept in full medium at 37C°C and 5% CO_2_, and pictures were acquired within a single z-plane that was set at the beginning by focusing on septin fibers of pre-mitotic cells, and kept by an autofocus system. NRK49F SEPT2-eGFP cells and gels for U-ExM were imaged with an Olympus spinning disk microscope (Yokogawa CSU-X1, Hamamatsu C11440 camera), using a 60x immersion oil objective (1.42 NA). Cells depicted in Fig. 2D-E were imaged within a single z-plane by focusing on septin fibers. For cells depicted in Fig. 2F, a z-stack with a spacing of 0.3 µm was acquired. For U-ExM, 20 images were acquired per condition, per experiment. For each image, a z-stack of 21 layers with a spacing of 1μm was acquired.

Image processing and analysis was performed with the open-source software Fiji (Image J).

Quantifications depicted in Fig. 1B-E were performed by identifying and manually counting mitotic cells displaying a mitotic spindle, or a cytokinetic bridge as schematized in fig. 1A. This number was then divided by the total number of nuclei, identified via a macro and serving as the total number of cells in the picture.

The qualitative assessment of anillin at the ICB was conducted on maximum intensity projections of z-stacks. For measuring the intensity of the acetylated tubulin bridge, average intensity projections of the z-stacks were generated, and the acetylated tubulin channel was segmented in order to obtain regions of interest (ROI) outlining the acetylated tubulin bridges. Subsequently, the fluorescence intensity of acetylated tubulin was measured as integrated density within the obtained ROI on average intensity z-projections, after background subtraction. The intensity of PRC1 bridges, MKLP1 and MgcRacGAP dots were measured the same way, in regular confocal or expanded samples. The length of the acetylated tubulin bridges was measured as the max Feret diameter of the ROI outlining the acetylated tubulin bridges. Septin enrichment at the acetylated tubulin bridge was determined by dividing the intensity of SEPT2 in a ROI outlining the acetylated tubulin bridge by the intensity of SEPT2 in a ROI outlining the whole dividing cell (drawn by hand); here as well measurements were carried out on average intensity z-projections, after background subtraction. In quantifications depicted in Fig. S1E/F, the intensity of PI(4,5)P_2_ per cell area was measured as mean grey value of the EGFP-PH-PLCδ1 fluorescence in the ROI outlining single cells (drawn by hand), after background subtraction. Intensity line scan analysis as depicted in Fig. 2F-G was performed on maximum intensity projections of the z-stacks. Intensity line scan analysis as depicted in Fig. S5C-D was performed on images acquired within a z-plane at the middle of the bridge.

For all displayed images brightness and contrast were adjusted equally for different conditions, unless otherwise indicated.

### Ultrastructure expansion microscopy (U-ExM)

Sample were processed using the U-ExM method^45^. Abbreviations used: FA (Formaldehyde), AA (Acrylamide), BIS (N,N’-Methylenebisacrylamide), TEMED (N,N,N′,N′-Tetramethylethylenediamine), APS (Ammonium persulfate), SA (Sodium acrylate). HeLa cells (seeded on 18 mm matrigel coated coverslips) were rinsed with PHEM buffer (60 mM PIPES, 25 mM HEPES, 2 mM MgCl_2_, 10 mM EGTA, pH 6.9), then fixed in 4% PFA, 0.1% Triton X-100 in PHEM at 37°C for 15 min. Fixed coverslips were rinsed in PHEM buffer again and transferred into 1 mL of anchoring solution (0.7% FA, 1% AA in PBS). Samples were incubated for 16 h at RT. U-ExM monomer solution (19% (wt/wt) SA, 10% (wt/wt) AA, 0.1% (wt/wt) BIS in PBS) was prepared one day before the gelation and stored at −20°C until use. 90 µL of monomer solution were mixed with 5 µL of 10% TEMED and 5 µL of 10% APS (0.5% APS and TEMED in final monomer solution) and placed on Parafilm in a drop in a precooled humid chamber. The coverslips were placed on the drops facing downwards and incubated on ice for 5 min. For gelation coverslips were incubated at 37°C for one hour in the humid chamber. Gels (still on coverslips) were placed in 2 mL denaturing buffer (200 mM SDS, 200 mM NaCl, 50 mM Tris, pH9) in a six well plate for 15 min at RT with agitation. For denaturation gels were transferred into 15 ml centrifuge tubes filled with 3 mL of denaturation buffer and incubated at 95°C in for one hour. Subsequently, the gels were washed several times with MilliQ water and then expanded in 10 cm dishes in MilliQ water. The water was exchanged twice after 30 min. Finally, gels were left to expand in water over night at 4°C.

Gels were measured with a caliper the following day to calculate the expansion factor. Gels were cut into 2×2 cm squares, and the squares were incubated twice in PBS for 15 min. The shrunken gels were stained with antibodies diluted in 2% BSA/PBS. To this end each gel piece was placed in a 12–well plate, covered with 500 µL of antibody solution and incubated for three hours at 37°C and at 120 rpm. Gels were then washed three times for 20 min with PBS-T (PBS supplemented with 0.1% Tween20) in a 6-well plate with gentle agitation. Secondary antibody incubation was executed as indicated above for 2.5 h. Gels were then washed three times for 20 min with PBS-T in a 6-well plate under gentle agitation. Final expansion was done in MilliQ water in 10 cm dishes, exchanging the water at least two times after 30 min, and letting the expansion plateau by an overnight incubation at 4°C. For imaging, gels were mounted in imaging dishes (zell-kontakt) with 2% agarose gels. To calculate the scale, the pixel size of the camera was divided by the measured gel expansion factor (∼4,2).

### Cell lysates

Cells were trypsinized, washed with PBS, and lysed for 15 minutes on ice in lysis buffer (20 mM HEPES, 100 mM KCl, 2 mM MgCl_2_, 1% Triton X-100, pH 7,4) supplemented with protease inhibitor cocktail (Sigma-Aldrich) and 1 mM PMSF. Lysates were cleared by centrifugation at 17000g for 15 minutes at 4°C, and the protein concentration was determined with Bradford reagent (Sigma-Aldrich). Lysates were subsequently denatured in Laemmli sample buffer for 5 minutes at 95°C. For each sample, 10 – 30 μg of protein were resolved by SDS-PAGE and analyzed by western blot (see below).

### Immunoprecipitation assay

HEK-293T cells seeded in 10 cm petri dishes were transfected at 70% confluency with 5 μg of plasmid encoding mCherry-PIPKIγ or mCherry alone and 5 μg of myc-tagged SEPT5, SEPT6, or SEPT9, using the calcium phosphate method. 24 h later cells were trypsinized, washed once with PBS and lysed at 4°C for 15 minutes in lysis buffer (20 mM HEPES, 100 mM KCl, 2 mM MgCl_2_, 1% Triton X-100, pH 7,4) supplemented with protease inhibitor cocktail (Sigma-Aldrich), 1 mM PMSF and phosphatase inhibitors cocktails 2 and 3 (Sigma-Aldrich). Lysates were cleared by centrifugation at 17000 x *g* for 15 minutes. 0.6 mL of the resulting supernatant (containing 2 – 4 mg of proteins) were supplemented with 15 μL of RFP-Trap magnetic particles (ChromoTek), and incubated for 2.5 h at 4°C on a rotating wheel. Beads were washed three times with lysis buffer, and a fourth time with lysis buffer without detergent. Proteins bound to the beads were eluted by boiling in 60 μL of Laemmli sample buffer for 5 minutes, resolved by SDS-PAGE and analyzed via immunoblot.

For immunoprecipitation experiments cell lysates derived from HeLa cells arrested in S-phase by thymidine treatment (obtained upon trypsinization), or from HeLa cells in cytokinesis (obtained by gentle pipetting) were incubated with 5 µg of rabbit-anti-human control or of rabbit-anti-SEPT2 antibodies, bound to protein A/G magnetic beads (Pierce^TM^) at 4°C for 3 – 4 h on a rotating wheel. Beads were washed four times in lysis buffer (20 mM HEPES, 100 mM KCl, 2 mM MgCl_2_, 1% Triton X-100, pH 7,4) and once in lysis buffer devoid of detergent. Retained proteins were released by boiling in 90 µL of Laemmli sample buffer for 5 minutes, resolved by SDS-PAGE and analyzed by Western blotting.

### GST pulldown assay

Expression and purification of GST-tagged PIPKIγ tails (or of GST alone, used as control) was performed essentially as described previously^59^. For pulldown experiments mouse brain extracts were obtained by homogenizing four brains in homogenization buffer (20 mM HEPES, 320 mM sucrose, pH 7.5) containing complete EDTA-free protease inhibitor cocktail (Roche). The homogenate was centrifuged at 1000 x *g* for 15 min at 4°C. The supernatant was recovered, supplemented with 1% Triton X-100, 100 mM KCl, 2 mM MgCl2, and incubated on ice for 10 min. The lysate was cleared by centrifugation at 17,000 x *g* for 15 min, and subsequently at 178,000 x *g* for 15 min (4°C). The supernatant was recovered and used at a concentration of 14 mg protein/mL. Pulldown experiments were performed by incubating 1 mL of protein extract with 70 μg of GST-fusion protein or of GST for 3h at 4°C by end-over-end rotation. The samples were subsequently washed four times with buffer containing 20 mM HEPES, 100 mM KCl, 2 mM MgCl2, 1% Triton X-100 pH 7.4, and once in the same buffer without detergent. Proteins were eluted from the beads by boiling for 5 minutes in 100 μL of Laemmli buffer, and analyzed by Western blotting. Chemiluminescent signals were detected using the ChemieDoc MP Imaging System (BioRad) controlled by the Image Lab 6.0.1 software. Fluorescent signals were detected using the LI-COR Odyssey Fc imaging system controlled by the Image Studio software.

### Analysis of the expression of PIPKIγ isoforms

RT-PCRs were performed as described previously^60^. Briefly, 1 µg of RNA was used with isoform-specific reverse primers for the RT-reaction, and the subsequent PCR was performed with a ^32^P-labeled forward primer. Products were separated by denaturing PAGE and quantified using a Phosphoimager (Typhoon 9200, GE Healthcare) and ImageQuantTL software. The sequence of the primers used for PCR and the resulting sizes of the amplicons were as follows: Common PIPKIγ forward: GCGCCCGCCACCGACATCTAC; PIPKIγ-i1-3 reverse: CATCTCCCGAGCTCTGGGCCTC (i1=125 nt, i2=210 nt, i3=290 nt); PIPKIγ-i4 reverse: GAGACCAGGACGCGCACAAACCAG (i4 = 154 nt); PIPKIγ-i5 reverse: CAGACACTGAGCTTCCGGCCGG (v5 = 195 nt)

### Statistics and reproducibility

All data were derived from at least three independent experiments and are presented as means ± standard deviation (SD). GraphPad Prism version 9.2 software was used for statistical analysis. Unpaired two-tailed t-test was applied to compare two groups. One-way ANOVA followed by Dunnett’s multiple comparison test was used to compare more than one experimental group to a control group. When the control group was set to 1 by normalization, one sample two-tailed t-test was applied for comparing one or more experimental groups to control. The level of significance is indicated in the figures by asterisks (*P < 0.05; **P < 0.01; ***P < 0.001; ****P < 0.0001), and detailed in the figure legends as exact P value.

### Antibodies used in this study

**Table 1:** Primary antibodies.

| Antibody | Source | Identifier | IF | U-exM | WB |
| --- | --- | --- | --- | --- | --- |
| Acetylated tubulin (mouse) | Sigma-Aldrich | T7451 | 1:2000 |  |  |
| α-tubulin (mouse) | Sigma-Aldrich | T5168 | | 1:400 | 1:2000 |
| β-tubulin (mouse) | Sigma-Aldrich | T5293 | | 1:400 | |
| PIPKIα (mouse) | Santa-Cruz | sc-398687 | | | 1:100 |
| PIPKIβ (mouse) | Santa-Cruz | sc-514169 | | | 1:100 |
| PIPKIγ (rabbit) | Home-made in this study | | | | 1:500 |
| SEPT2 (rabbit) | Sigma-Aldrich | HPA018481 | 1:200 | 1:200 | 1:500 |
| SEPT5 (mouse) | Santa-Cruz | sc-20040 |  |  | 1:100 |
| SEPT6 (rabbit) | Home-made in this study |  | 1:70 |  | 1:250 |
| SEPT7 (rabbit) | Santa-Cruz | sc-20620 | 1:100 |  | 1:500 |
| SEPT7 (rabbit) | TECAN | JP18991 | 1:250 | 1:200 |  |
| SEPT3 (mouse) | Sigma-Aldrich | WH0055964M3 |  |  | 1:1000 |
| SEPT9 (rabbit) | (Diesenberg et al., 2015) |  | 1:400 |  |  |
| Talin (mouse) | Sigma-Aldrich | T3287 |  |  | 1:1000 |
| GAPDH (mouse) | Sigma-Aldrich | G8795 |  |  | 1:10000 |
| Anillin (goat) | Abcam | ab5910 | 1:50 |  |  |
| c-Myc (mouse) | Hybridoma clone obtained from DSHB, purified ourselves | 9E10 |  |  | 1:400 |
| RFP (rabbit) | Clontech | 632496 |  |  | 1:1000 |
| HA (mouse) | Abcam | ab18181 | 1:500 |  |  |
| OCRL1 (rabbit) | Cell Signaling | 8797 |  |  | 1:500 |
| PRC1 (mouse) | Thermo-Fisher | MA1-846 | 1:1000 |  |  |
| CIT-K (mouse) | BD Transduction Laboratories | 611377 | 1:300 |  | 1:500 |
| MKLP1 (rabbit) | GeneTex | GTX120875 | 1:250 | 1:250 | 1:500 |
| MgcRacGAP (goat) | Abcam | Ab2270 |  | 1:200 |  |
| MgcRacGAP (rabbit) | Proteintech | 13739-1-AP | 1:500 |  | 1:500 |
| GFP (mouse) | Clontech | 632381 |  |  | 1:1000 |
| GFP (rabbit) | Abcam | ab6556 | 1:1000 |  |  |
| PI(4,5)P<sub>2</sub> (mouse, IgM) | Echelon Biosciences | Z-A045 | 1:400 |  |  |

**Table 2:** Secondary antibodies.

| Antibody | Conjugate | Source | Identifier | IF | U-exM | WB |
| --- | --- | --- | --- | --- | --- | --- |
| Goat anti rabbit | Alexa Fluor 488 | Thermo-Fisher | A-11034 | 1:200<br>1:400<br>(PH-PLC staining) |  |  |
| Goat anti mouse | Alexa Fluor 568 | Thermo-Fisher | A-11004 | 1:200 |  |  |
| Donkey anti goat | Alexa Fluor 488 | Thermo-Fisher | A-11055 | 1:200 | 1:250 |  |
| Donkey anti rabbit | Alexa Fluor 647 | Thermo-Fisher | A-31573 | 1:200 | 1:250 |  |
| Donkey anti mouse | Alexa Fluor 568 | Thermo-Fisher | A-10037 | 1:200 | 1:250 |  |
| Goat anti mouse | Alexa Fluor 647 | Thermo-Fisher | A21236 | 1:200 |  |  |
| Goat anti mouse (IgM) | Alexa Fluor 568 | Thermo-Fisher | A21043 | 1:500 |  |  |
| Goat anti mouse | HRP | Jackson ImmunoResearch | 115-035-003 |  |  | 1:2500 |
| Goat anti rabbit | HRP | Jackson ImmunoResearch | 111-035-003 |  |  | 1:2500 |
| Goat anti mouse | IRDye® 800CW | LI-COR | 926-32210 |  |  | 1:5000 |
| Goat anti rabbit | IRDye® 680RD | LI-COR | 926-68071 |  |  | 1:5000 |

