## Supplemental Figures for "Septin-associated PIPKIγ splice variants drive centralspindlin association with the midbody via PI(4,5)P_2_"

### Supplemental Information

#### Supplemental Figures

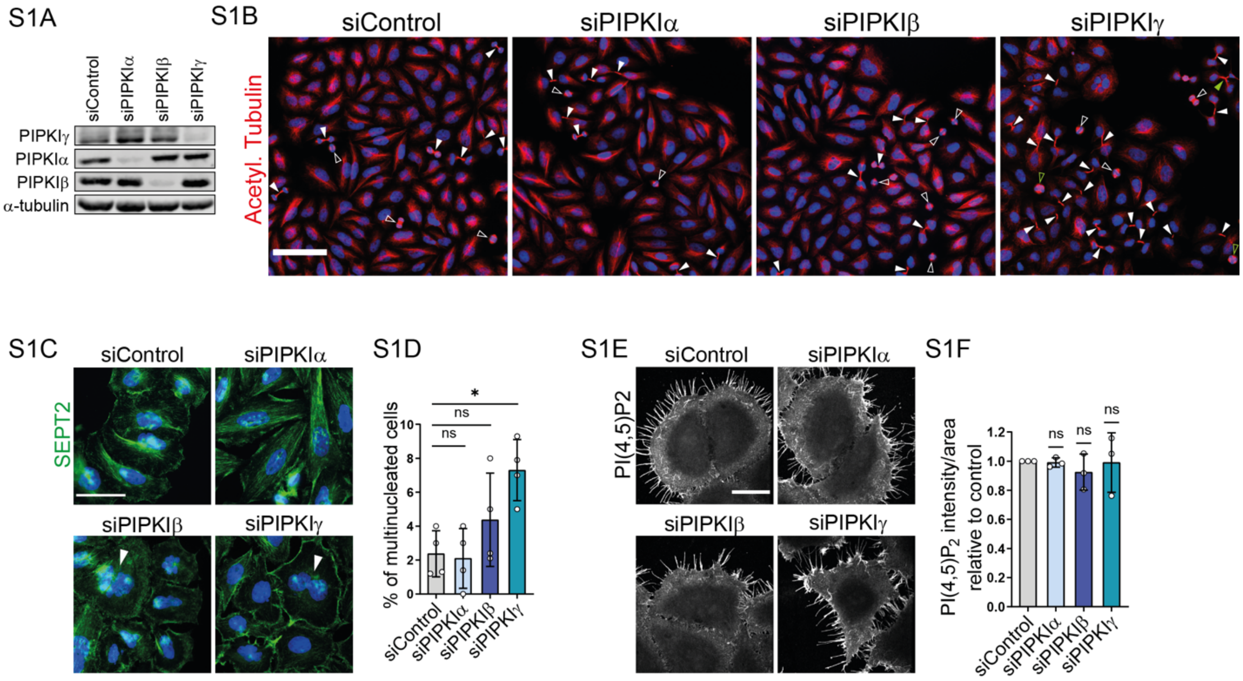


**Figure S1 (related to Figure 1) (A)** Western blot analysis show efficient knock-down of PIPKI isozymes. **(B)** HeLa were treated with siRNA control or against PIPKIα, β, γ, and stained for acetylated tubulin and DAPI. Four areas of 1,3 mm$^2$ (between 2593-4431 cells) were imaged by semi-automated imaging (epi-fluorescence) per coverslip per experiment (see Fig.1A-E for quantifications). Representative insets are shown. White arrowheads indicate acetylated tubulin spindles (open), and acetylated tubulin bridges (filled), green arrowheads indicate multipolar spindles (open) and multipolar bridges (filled). Scale bar: 100µm. **(C-D)** Depletion of PIPKIγ increases multinucleation. **(C)** Representative epi-fluorescent images of HeLa cells treated with control siRNA, or with siRNA targeting PIPKIα, β, or γ, and immunostained for SEPT2 and with DAPI, scale bar: 50µm. **(D)** Fraction of multinucleated cells. Data are depicted as mean ± SD (n=4; ~ 200 cells were imaged per condition per experiment). Statistics: 1-way ANOVA, followed by Dunnett's multiple comparison test. P-values: siControl vs. siPIPKIα, P=0.9944; siControl vs. siPIPKIβ, P=0.3827; siControl vs. siPIPKIγ, P=0.0113. **(E-F)**, Depletion of PIPKIα, β, or γ does not cause major changes in plasmalemmal $PI(4,5)P_2$. **(E)** Representative confocal images of HeLa cells treated with control siRNA, or with siRNA targeting PIPKIα, β, or γ, and stained for $PI(4,5)P_2$ with purified EGFP-tagged PH-PLC δ1 domain, scale bar: 30µm. **(F)** Relative intensities of $PI(4,5)P_2$ per cell area, depicted as mean ± SD (n=3; ~ 30 cells were imaged per condition per experiment). Statistics: two-tailed one sample t test (hypothetical value: 1). P-values: siPIPKIα, P=0.6580; siPIPKIβ, P=0.4034; siPIPKIγ, P=0.9415.

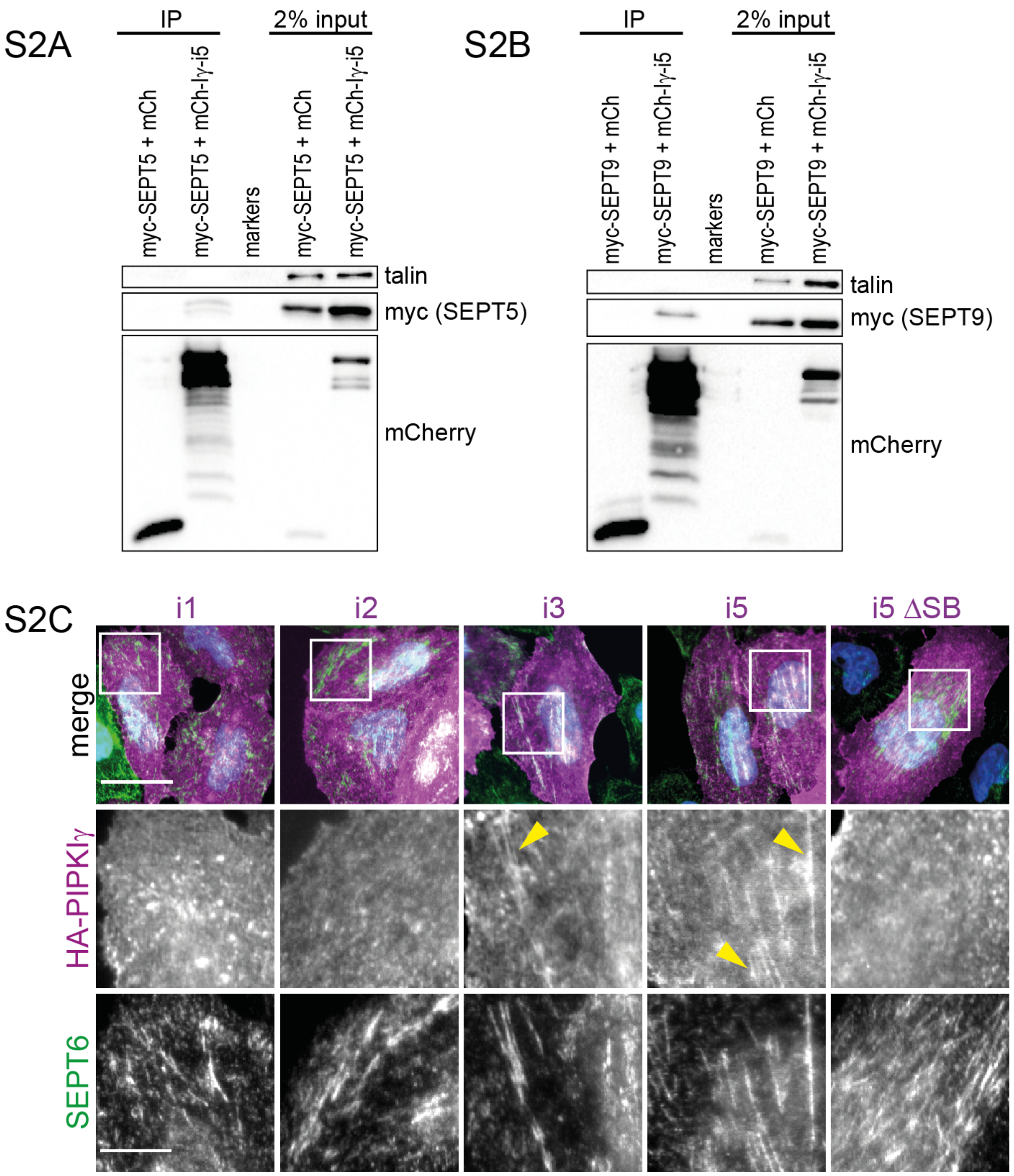


**Figure S2 (related to Figure 2) (A-B)** Co-immunoprecipitation of myc-tagged SEPT5 **(A)** or SEPT9 **(B)** with mCherry-tagged PIPKIγ-i5 from HEK-293T cell lysates. **(C)** Representative epi-fluorescent images derived from transfected HeLa cells. Over-expressed HA-tagged PIPKIγ-i3/i5, but not i1/i2, or i5ΔSB, exhibit a filamentous pattern (yellow arrowheads) overlapping with endogenous septin filaments (revealed by immunostaining of SEPT6) in HeLa cells. Scale bar, 30 µm (merge), and 10 µm (insets).

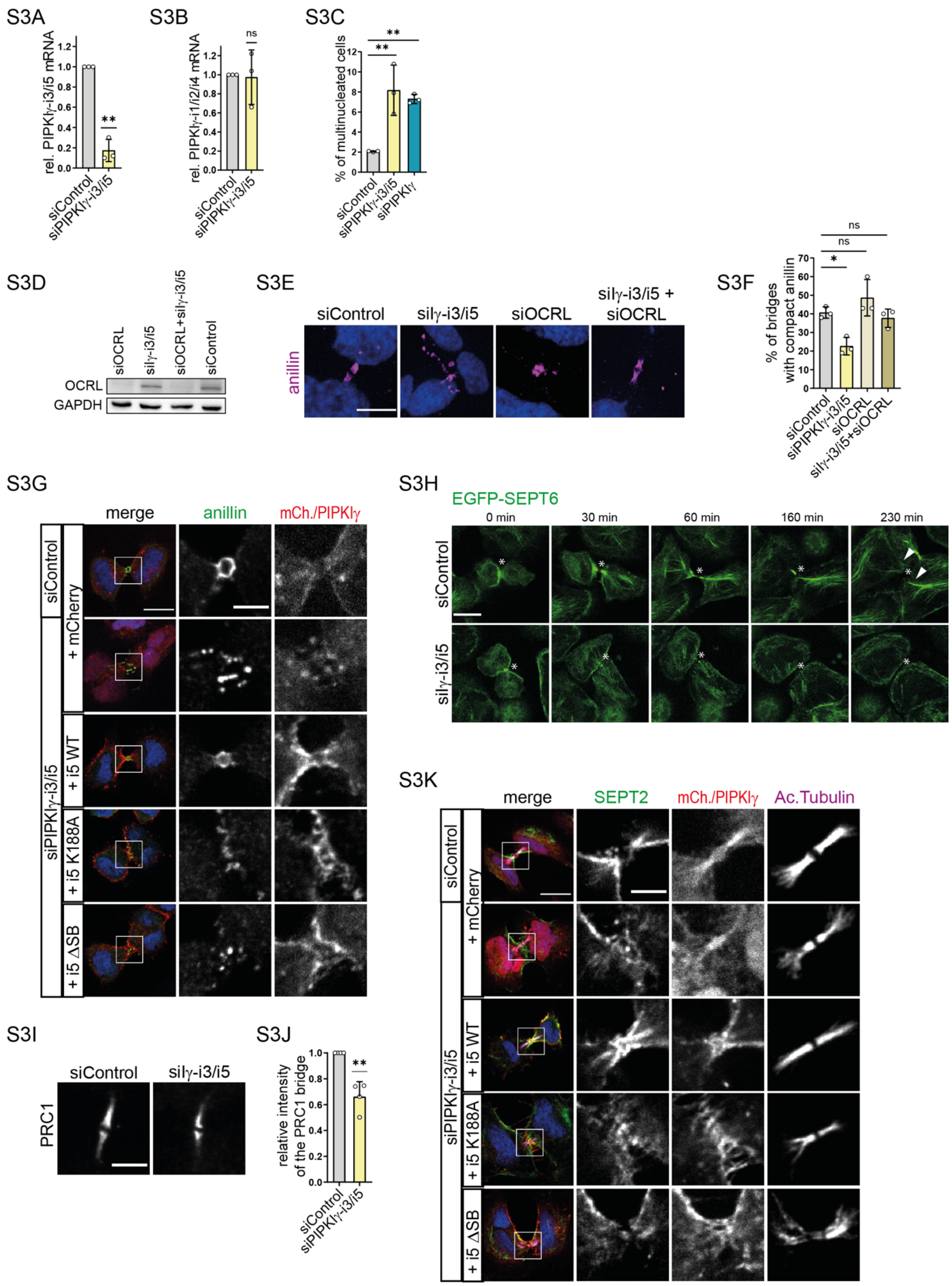


**Figure S3 (related to Figure 3) (A-B)** Knockdown of PIPKIγ-i3/i5 is specific and efficient. **(A)** Expression of PIPKIγ-i3+i5, normalized to GAPDH, and relative to control. **(B)** Expression of PIPKIγ-i1+i2+i4, normalized to GAPDH, and relative to control. Data are depicted as mean ± SD (n=3). Statistics: two-tailed, one sample t-test (hypothetical value of 1). P values: P= 0,0059 (A) and P= 0,8865 (B). **(C)** Fraction of multinucleated cells upon treatment with isoform-specific siRNA, or with siRNA targeting all

PIPKIγ isoforms. Data are depicted as mean ± SD (n=4; ~ 200 cells were imaged per condition per experiment). Statistics: 1-way ANOVA, followed by Dunnett's multiple comparison test. Adjusted P values: siControl vs. siPIPKIγ-i3/i5, P=0.0041; siControl vs. siPIPKIγ, P=0.0086. **(D-F)** Co-depletion of OCRL and PIPKIγ-i3/i5 rescues anillin accumulation at the midbody. **(D)** Western blot analysis of lysates derived from HeLa cells treated with indicated siRNAs. **(E)** Representative confocal pictures (maximal intensity z-projections) of anillin at ICBs of HeLa cells upon depletion of PIPKIγ-i3/i5, of OCRL, or of both. Cells were synchronized at late stages of cytokinesis, and immunostained for anillin. Scale bar, 10μm. **(F)** Fraction of ICBs with compact anillin. Values indicate mean ± SD (n=3; ~ 30 cytokinetic cells per condition per experiment were imaged). Statistics: 1-way ANOVA, followed by Dunnett's multiple comparison test. Adjusted P values: siControl vs siPIPKIγ-i3/i5, P=0.0182; siControl vs. siOCRL, P=0.3236; siControl vs siPIPKIγ-i3/i5 + OCRL, P=0.8808. **(G)** Representative confocal images derived from rescue experiments. Cells were treated with control or PIPKIγ-i3/i5-targeting siRNA, and transfected with mCherry, or mCherry (mCh.)-tagged PIPKIγ-i3/i5 variants. Synchronized cells were fixed at late stage of cytokinesis, and immunostained for anillin. Scale bars, 15 μm (merge), 5μm (insets). **(H)** Depletion of PIPKIγ-i3/i5 impairs septin organization at the ICB. Genome-edited HeLa cells expressing EGFP-SEPT6 cells were synchronized, and imaged throughout cytokinesis by confocal microscopy. Images show representative frames derived from movies of control- or PIPKIγ-i3/i5-depleted cells. Scale bar, 15μm. Asterisk indicates the putative midbody; arrowheads indicate sinuous septin fibers possibly deriving from the cytokinetic bridge. **(I-J)** Depletion of PIPKIγ-i3/i5 reduces PRC1 levels at the cytokinetic bridge. **(I)** Representative confocal images (max intensity z-projections) of PRC1 at the ICB, derived from control, or from PIPKIγ-i3/i5-depleted HeLa cells, upon synchronization at late stages of cytokinesis and immunostaining. Scale bar, 5 μm. **(J)** Relative intensity of PRC1. Quantification was performed on average intensity z-projections after background subtraction. Normalized data are depicted as mean ± SD (n=4; 15-30 cytokinetic cells were imaged per condition per experiment). Statistics: two-tailed one sample t test (hypothetical value: 1; P: 0.0097). **(K)** Representative confocal images derived from rescue experiments. Cells were treated with control or PIPKIγ-i3/i5-targeting siRNA, and transfected with mCherry, or mCherry (mCh.)-tagged PIPKIγ-i3/i5 variants. Synchronized cells were fixed at late stage of cytokinesis, and immunostained for acetylated tubulin and SEPT2. Scale bars, 15 μm (merge), 5 μm (insets).

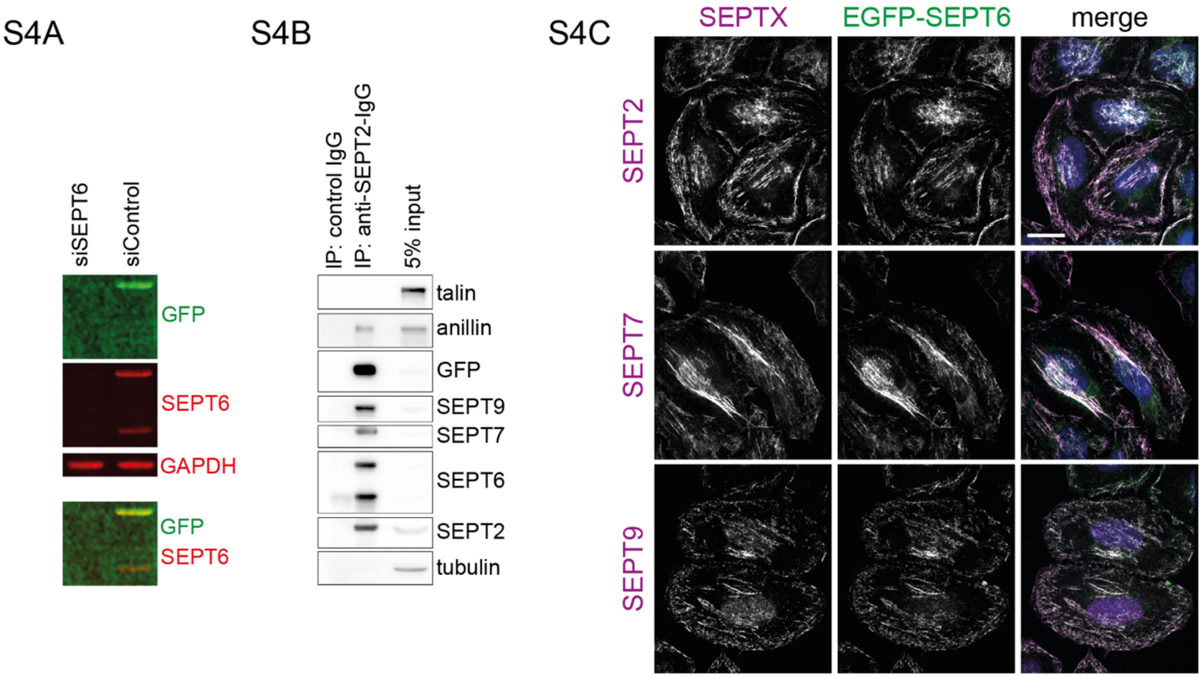


**Figure S4 (related to Figure 3) (A-C)** Characterization of genome-edited HeLa cells expressing EGFP-SEPT6. EGFP-SEPT6 interacts with other septins and is incorporated into septin fibers. **(A)** Western blot analysis of lysates derived from genome-edited, EGFP-SEPT6-expressing HeLa cells treated with SEPT6-specific or control siRNAs. Lysates were separated by SDS-PAGE, and immunoblotted for GFP, SEPT6 or GAPDH. **(B)** Western blot analysis of SEPT2-immunoprecipitates obtained from lysates of genome-edited, EGFP-SEPT6 expressing HeLa cells. Precipitates were separated by SDS-PAGE, and immunoblotted for the indicated proteins. SEPT2 efficiently co-precipitates EGFP-SEPT6 and SEPT6, as well as other septin paralogs and anillin. **(C)** Representative confocal images derived from genome-edited, EGFP-SEPT6 expressing HeLa cells. Cells were fixed, and immunostained for endogenous SEPT2, SEPT7, or SEPT9. EGFP-SEPT6 is efficiently incorporated into septin filaments. Scale bar, 10 µm.

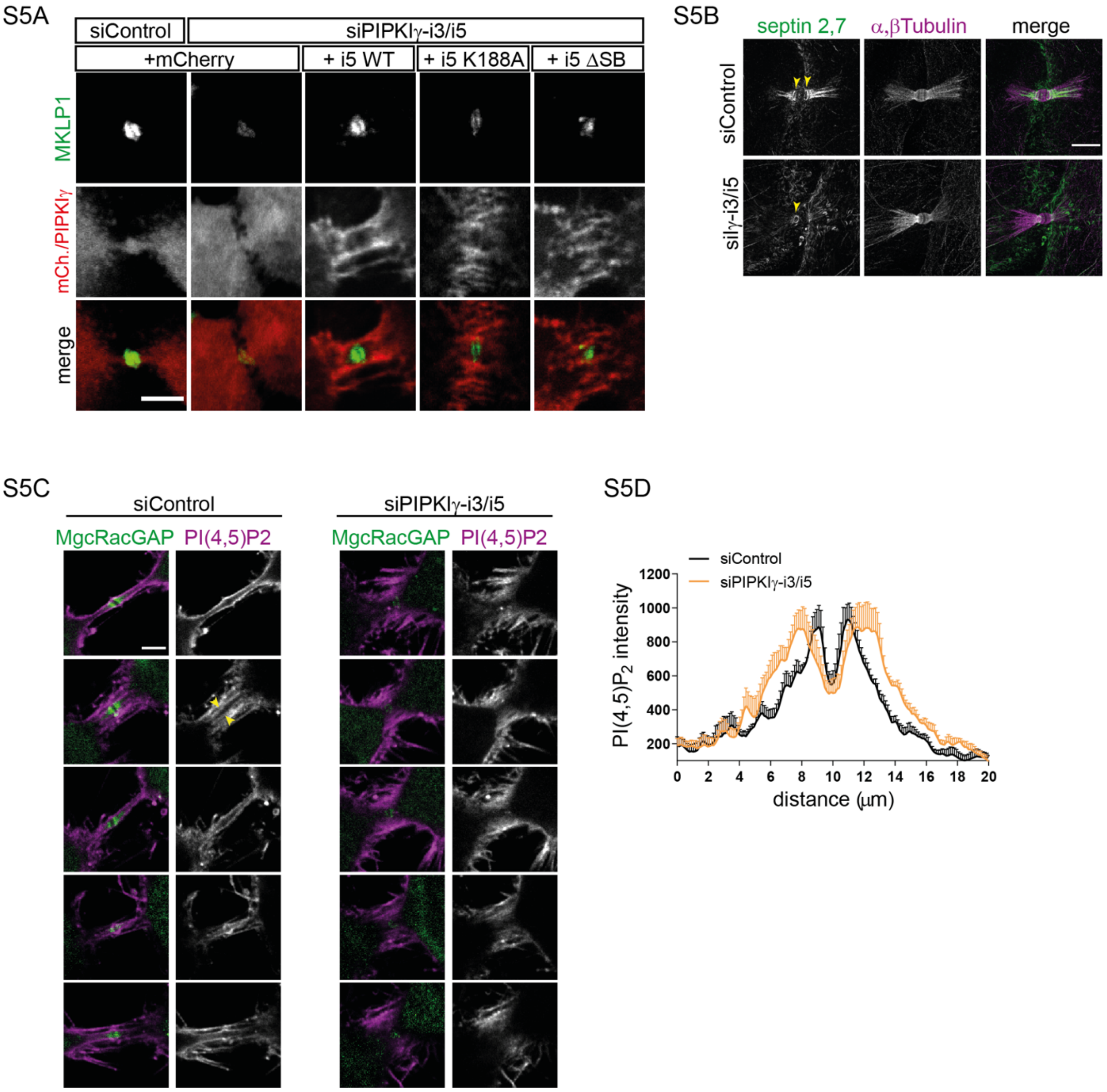


**Figure S5 (related to Figure 4) (A)** Representative confocal images derived from rescue experiments. Cells were treated with control or PIPKIγ-i3/i5-targeting siRNA, and transfected with mCherry, or mCherry (mCh.)-tagged PIPKIγ-i3/i5 variants. Synchronized cells were fixed at late stage of cytokinesis, and immunostained for MKLP1. Scale bar, 5μm. **(B)** Confocal images derived from U-ExM indicate redistribution of septins away from bridge microtubules in the absence of PIPKIγ-i3-i5. Cells were immunostained for SEPT2/SEPT7 and α-/β-tubulin, and imaged on a spinning disk confocal microscope. Representative images (max-intensity projections of 21 slices with 1 μm spacing) are shown. Scale bar, 10 μm. **(C-D)** Depletion of PIPKIγ-i3/i5 alters the distribution of $PI(4,5)P_2$ across the intercellular bridge. **(C)** Representative confocal images of HeLa cells upon treatment with control siRNA, or siRNA targeting PIPKIγ-i3/i5. Cells were synchronized at late stages of cytokinesis, and stained for $PI(4,5)P_2$ and MgcRacGAP. Scale bar, 5 μm. **(D)** $PI(4,5)P_2$ intensity profile along a 20 μm line centered in the middle of each bridge (supposed localization of the midbody). Data are represented as mean ± SEM (n=5, 15-30 bridges were imaged per condition per experiment).
